# Carbomer-based Nano-Emulsion Adjuvant Enhances Dendritic Cell Cross-presentation via Lipid Body Formation Independent of Glycolysis

**DOI:** 10.1101/2020.05.08.083790

**Authors:** Woojong Lee, Brock Kingstad-Bakke, Brett Paulson, Autumn R. Larsen, Katherine Overmyer, Chandranaik B. Marinaik, Kelly Dulli, Randall Toy, Gabriela Vogel, Katherine P. Mueller, Kelsey Tweed, Alex J. Walsh, Jason Russell, Krishanu Saha, Leticia Reyes, Melissa C. Skala, John-Demian Sauer, Dmitry M. Shayakhmetov, Joshua Coon, Krishnendu Roy, M. Suresh

## Abstract

Here, we report that a carbomer-based adjuvant, Adjuplex® (ADJ), stimulated robust CD8 T-cell responses to subunit antigens by modulating multiple steps in the cytosolic pathway of cross-presentation, and afforded effective immunity against virus and intracellular bacteria. Cross-presentation induced by TLR agonists requires a critical switch to anabolic metabolism, but ADJ enhanced cross presentation without this metabolic switch in DCs and NLRP3-driven caspase 1 activity. Instead, ADJ induced in DCs, an unique metabolic state, typified by dampened oxidative phosphorylation and basal levels of glycolysis. In the absence of increased glycolytic flux, induction of ROS and lipid bodies (LBs) and alterations in LB composition mediated by ADJ were critical for DC cross-presentation. These findings challenge the prevailing metabolic paradigm by suggesting that DCs can perform effective DC cross-presentation, independent of glycolysis to induce robust T cell-dependent protective immunity to intracellular pathogens. These findings have implications in the rational development of novel adjuvants.

## INTRODUCTION

Development of effective vaccines remains the central principle for controlling infectious diseases in humans and animals. Typically, vaccines can be classified into the following categories: replicating vaccines (live-attenuated viruses; e.g. smallpox), and non-replicating vaccines (subunit [hepatitis B], virus-like particles [human papilloma virus], toxoid [tetanus], and conjugated vaccines [*Haemophilus influenzae* type B]) (Pulendran and Ahmed, 2011). To date, protection afforded by the most effective vaccines is primarily dependent upon the elicitation of antibodies (Burton, 2002). By contrast, development of vaccines against infections that requires T cells for pathogen control, such as HIV, tuberculosis (TB), and malaria, remains a difficult challenge for vaccinologists (Hoft, 2008, Reyes-Sandoval et al., 2009, Walker et al., 2011). Live-attenuated vaccines are highly immunogenic and elicit both humoral and cell-mediated immunity, but their use can be contraindicated during pregnancy and in immunocompromised individuals (Kitchener, 2004, Lindsey et al., 2016, Struchiner et al., 2004). Subunit or inactivated antigens are generally safe, but are poorly immunogenic unless formulated in pharmaceutical agents called adjuvants (Brito and O’Hagan, 2014).

CD8 T cell responses to non-replicating subunit protein antigens requires antigen cross-presentation by dendritic cells (DCs) (Bevan, 1976). Likewise, DC cross-presentation plays a pivotal role in eliciting CD8 T cell responses to intracellular pathogens (e.g. *Listeria monocytogenes*) and tumor antigens (Huang et al., 1994, Wolkers et al., 2001, Belz et al., 2005). Cross-presentation of MHC I-restricted antigens to CD8 T cells can occur via vacuolar or cytosolic pathways (Grotzke et al., 2017). In the vacuolar pathway, exogenous antigens are internalized into endosomes and digested by residential cathepsins (Shen et al., 2004). By contrast, in the cytosolic pathway, internalized antigens localize to the alkaline endosomal compartment, followed by antigen export into cytosol and downstream processing by the proteasomes. (Kovacsovics-Bankowski and Rock, 1995). Peptides resulting from proteasomal processing are translocated to endoplasmic reticulum (ER) by TAP1 transporter or to endosomes, and are loaded on to MHC-I molecules (Song and Harding, 1996).

Apart from engaging the appropriate antigen processing cellular machinery, metabolic reprogramming of DCs is considered to be an important facet of effective cross-presentation and activation of naïve T cells (Pearce and Everts, 2015). DC activation by TLR agonists triggers a metabolic switch from catabolic metabolism to anabolic metabolism, to accommodate increasing cellular demands for executing cellular functions, such as production of pro-inflammatory cytokines, upregulation of co-stimulatory molecules, and directed migration to draining lymph nodes (Wculek et al., 2019). Hence, understanding of DC metabolism is crucial for rationally designed vaccines that can effectively induce robust CD8 T cell responses to subunit protein antigens via mechanisms of DC cross-presentation.

Currently, there are only seven FDA-approved adjuvants for human use, and vaccines based on these adjuvants have mainly been evaluated for elicitation of humoral immunity (Del Giudice et al., 2018). There is high level of interest in developing adjuvants that can stimulate potent CD8 and CD4 T cell responses to subunit antigens. Carbomer (acrylic acid polymers)-based adjuvants (CBA) are components of several veterinary vaccines, and known to safely elicit potent neutralizing antibodies to malarial and HIV envelope glycoproteins in mice and non-human primates (Menon et al., 2015, Gupta et al., 2014, Anlar et al., 1993). We have also reported that the carbomer-based nano-emulsion adjuvant, Adjuplex® (ADJ; Advanced Bioadjuvants), elicits tissue-resident memory CD8 T cells in the lungs, and protects against influenza A virus in mice (Gasper et al., 2016). However, the mechanisms underlying the stimulation of protective CD8 T cells immunity by ADJ are largely unknown. In this manuscript, by employing integrative multidisciplinary approaches, we have systematically explored the mechanisms underpinning the molecular and metabolic basis for the potent activation of CD8 T cell immunity by ADJ.

## RESULTS

### Carbomer-based nano-emulsion adjuvant ADJ enhances cross-presentation of antigens by DCs in vitro *and* in vivo

We had previously reported that ADJ, a nano-emusion adjuvant was effective in eliciting CD8 T cell-dependent immunity to influenza A virus (Gasper et al., 2016). Here, we investigated mechanism of action of ADJ in driving antigen cross-presentation to T cells by DCs. We assessed whether exposure of bone marrow-derived DCs (BMDCs) to ADJ leads to enhanced antigen cross-presentation to CD8 T cells *in vitro* and *in vivo*. To qualitatively assess the magnitude of ADJ-mediated cross presentation of OVA antigen to CD8 T cells *in vitro*, BMDCs were treated with ADJ+OVA or OVA alone, and then were evaluated for their capacity to activate SIINFEKL-specific B3Z T cell hybridoma cells using a reporter assay (Karttunen et al., 1992). Here, we found that DCs stimulated with ADJ+OVA significantly induced β-gal in B3Z cells compared to OVA only control, suggesting enhanced antigen cross-presentation by ADJ-treated DCs (**Fig. 2A**). Next, to assess whether DCs treated with ADJ possess enhanced cross-priming abilities *in vivo*, we adoptively transferred DCs pretreated *in vitro* with ADJ+OVA, LPS+OVA, or OVA alone into C57BL/6 mice. The percentages and total numbers of SIINFEKL-specific CD8 T cells were significantly higher in spleens of mice that received DCs treated with ADJ+OVA, as compared to mice that received DCs treated with LPS+OVA or OVA only (**Fig. 2B**). In summary, data in **Fig. 1A-B** strongly suggested that ADJ enhances cross-presentation of antigens to CD8 T cells *in vitro* and *in vivo*.

**Figure 1.**
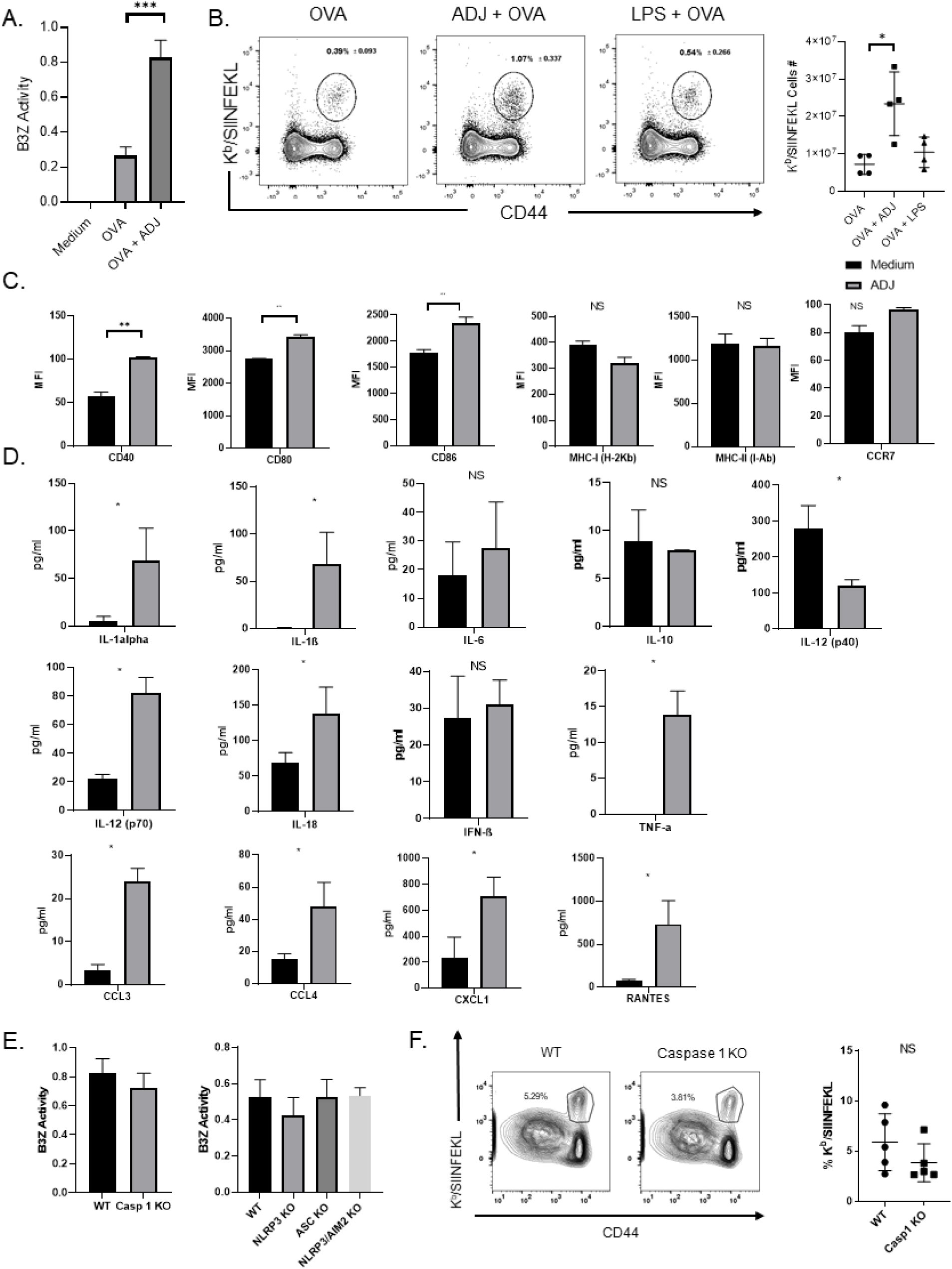
Carbomer-based adjuvants enhance DC cross-presentation, which is independent of co-stimulatory molecules and inflammasome activation. **(A)** BMDCs were exposed to media or OVA ± ADJ for 5 h, and co-cultured with B3Z cells for 24 h. β-gal in activated B3Z cells was quantified by CPRG colorimetry. (B) BMDCs were treated with OVA ± ADJ or LPS for 6 h, washed, and injected *i.v.* into C57BL/6 mice. After 7 days, CD8 T cells specific to the OVA SIINFEKL epitope were quantified using MHC I tetramers. Data are representative of ≥2 independent experiments. (C) FACS analysis of CD40, CD80, CD86, CCR7, MHC-I and MHC-II expression in BMDCs after treatment with ADJ for 6 h. (B) Measurement of cytokines in culture media of BMDCs 24 hr. after incubation with ADJ, by Multiplex Luminex Assay or ELISA. (D) β-gal production by B3Z cells after co-incubation with WT or DCs deficient for AIM2, ASC, NLRP3 or Caspase 1, pre-treated with OVA ± ADJ for 5 h. (E) Wild type and caspase 1 KO mice were vaccinated SQ with OVA ± ADJ. On the 8th day after vaccination, the percentages of activated OVA SIINFEKL-specific CD8 T cells in the spleen were quantified by flow cytometry. Data are representative of ≥2 independent experiments. Error bars show SEM; \**P*<0.01 (Student’s t-test and one-way ANOVA).

**Figure 2.**
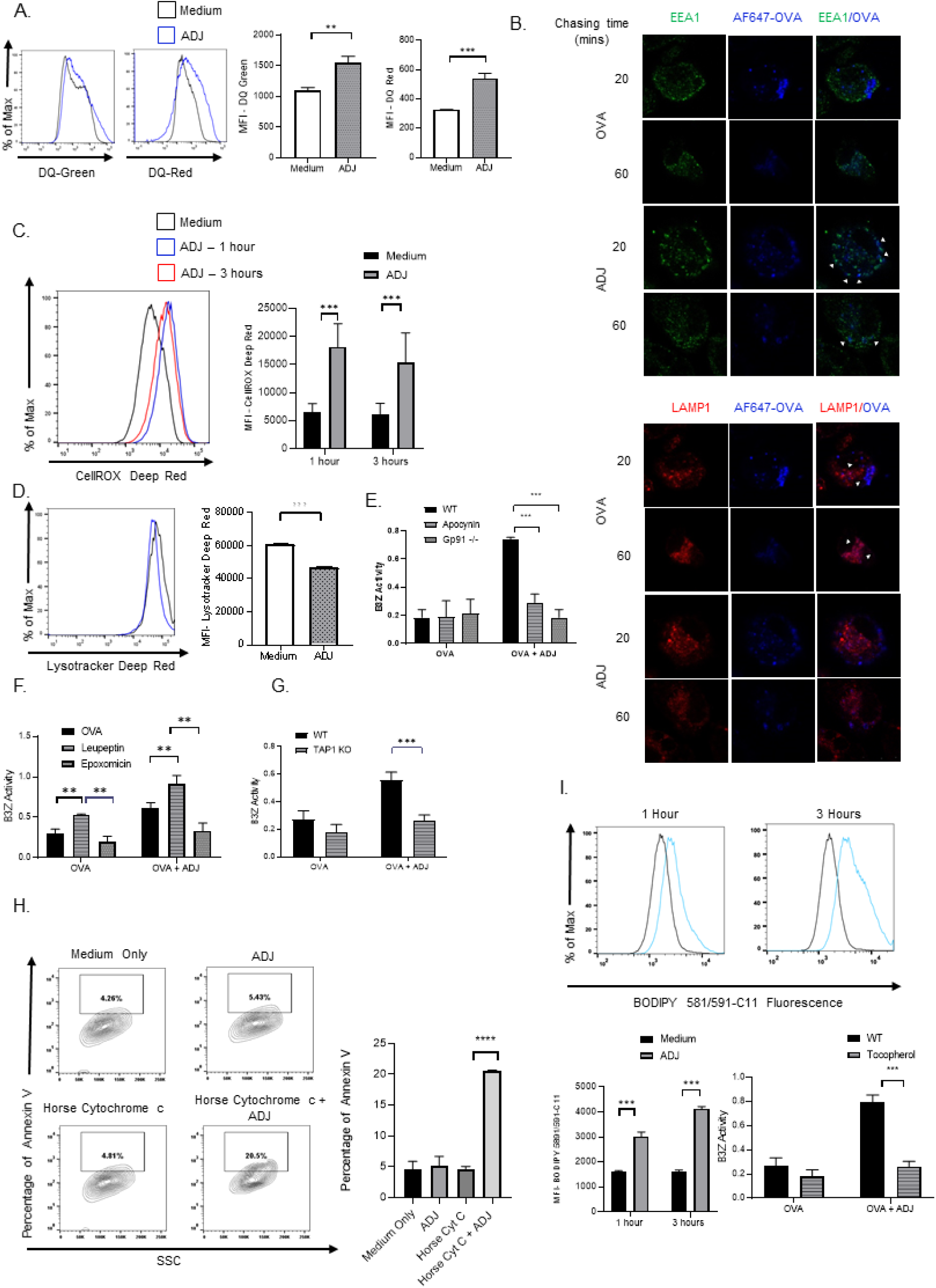
Carbomer-based adjuvant engages the endosome-to-cytosol pathway of cross-presentation. (A) BMDCs were cultured in DQ-OVA ± ADJ for 6 h and the fluorescence of DQ-green/red was measured using flow cytometry (B) BMDCs were pulsed with Alexa 647-OVA ± ADJ, chased at the indicated time-points, and immunostained to assess the degree of OVA co-localization with either EEA1^+ve^ (early endosomes) or LAMP1^+ve^ (lysosomes) organelles. Scale bars, 10 μm. (C-D) BMDCs were cultured in media ± ADJ for the indicated time and stained with CellRox and Lysotracker dyes to detect total cellular ROS and intracellular acidity, respectively. The fluorescence of CellROX and Lysotracker was measured using flow cytometry. (E) B3Z cross-presentation assay was performed using BMDCs deficient in gp91^-/-^ or DCs treated ± the NADPH oxidase inhibitor (apocynin). (F, I) β-gal production by B3Z cells after co-incubation with ADJ/OVA-treated BMDCs pre-treated with OVA ± ADJ in the presence/absence of tocopherol (I), leupeptin (F), or epoxomycin (F) for 5 h. (G) β-gal production by B3Z cells after co-incubation with ADJ/OVA-treated WT or DCs deficient for TAP1 pre-treated with OVA ± ADJ for 5 h. (H) ADJ-treated DCs were cultured with horse cytc ± ADJ to assess endosomal leakage of antigen into cytosol. Cell viability (as a read-out for antigen leakage) was measured by Annexin V staining. (I) BMDCs were cultured in media ± ADJ for 1 and 3 h, stained with BODIPY 581/591-C11, and a blue-shift of BODIPY 581/591-C11 fluorescence was measured using flow cytometry. Data are representative of ≥2 independent experiments. Error bars are the SEM; \*\**P*<0.001; \*\*\**P*<0.0001 (Student’s t-test and one-way ANOVA).

### ADJ induces inflammasome activation in DCs, but deficiency for NLRP3, ASC or caspase 1 did not affect ADJ-mediated cross-presentation

Next, we examined the effects of ADJ on the expression profiles of cytokines, chemokines and canonical cell surface markers of DC activation. Compared to untreated DCs, ADJ-treated DCs showed statistically significant, yet modest increases in expression of CD40, CD80, and CD86; no significant differences in expression were observed for MHC-I, MHC–II, or CCR7 (**Fig. 1C**). ADJ-treated DCs produced higher levels of IL-12 (p70), TNF-α, IL-1α, CCL3, CCL4, CXCL1, and RANTES, as compared to untreated DCs; no significant differences in expression were observed for IL-6, IL-10, and IFN-β (**Fig. 1D**). Particularly, ADJ-stimulated DCs also produced significantly elevated levels of IL-1β and IL-18 (**Fig. 1D**), which is suggestive of inflammasome activation. Since inflammasome activation has been implicated in modulation of antigen presentation by DCs (Sokolovska et al., 2013, Li et al., 2019), we performed B3Z assays using DCs deficient in NLRP3, ASC, or caspase 1 to interrogate whether inflammasome activation is required for ADJ-induced cross-presentation in BMDCs. Surprisingly, loss of NLRP3, ASC or caspase 1 activity did not affect ADJ-induced cross-presentation by DCs, *in vitro* (**Fig. 1E**). To validate whether caspase 1 is required for cross-presentation *in vivo*, we immunized cohorts of wild type (WT) and caspase 1-deficient mice with ADJ+OVA, and quantified OVA SIINFEKL-specific CD8 T cells in spleens using MHC I tetramers at day 8 after immunization. Consistent with our results from B3Z assays, caspase 1 deficiency did not significantly affect the activation of SIINFEKL-specific CD8 T cells in spleens (**Fig. 1F**), suggesting that caspase 1 is not essential for ADJ-driven cross-presentation to CD8 T cells in vivo.

### ADJ modulates antigen processing and subcellular localization in DCs

We next examined how ADJ affects the dynamics of antigen processing in DCs. First, we examined the effect of ADJ on antigen uptake in DCs by culturing with OVA that was labeled with pH-insensitive dye, Alexa Fluor 647. Interestingly, we found that antigen uptake was significantly reduced in DCs treated with ADJ (**Fig. S1**). This finding was not totally unexpected, as some TLR agonists are known to reduce uptake of soluble antigen, yet increase antigen cross-presentation. (Tirapu et al., 2009). Next, we examined whether ADJ affected antigen processing, by treating DCs with DQ-OVA, which emits green fluorescence upon proteolytic degradation and red fluorescence upon subsequent aggregation of digested peptides. We found that ADJ enhanced OVA degradation and/or accumulation of processed OVA, as indicated by an increase of both DQ-green and DQ-red fluorescence at 6 hours (**Fig. 2A**). Hence, these data suggest that ADJ might dampen antigen uptake, but enhances antigen processing and/or accumulation of processed antigen in DCs.

Apart from antigen degradation, trafficking of antigens to the appropriate less acidic vesicular compartment, such as early endosomes, is central for efficient cross-presentation (Burgdorf et al., 2007, Xia et al., 2015). Hence, we analyzed the effect of ADJ on the intracellular routing of antigen using pHAB-OVA, which emits green fluorescence in the acidic compartment. In ADJ-treated DCs, pHAB-OVA was routed to a more alkaline compartment in DCs within 30 minutes, as indicated by lower fluorescence of pHAB-OVA (**Fig. S1**). In order to precisely localize antigen in ADJ-treated DCs, we investigated the extent to which OVA antigen was localized to early endosomes or lysosomes, using confocal microscopy. Microscopic images showed that in ADJ-treated DCs, antigen was preferentially co-localized with early endosomes (EEA1), rather than lysosomes (LAMP1) (**Fig. 2B**). Unlike in ADJ-treated DCs, antigen was localized to the acidic lysosomes in DCs that were only treated with OVA (**Fig. 2B**). To further corroborate our findings, we employed a co-localization analysis of microscopy images using Pearson’s correlation coefficients (PCC) (Adler and Parmryd, 2010). This analysis showed that the PCC of EEA1 with OVA was higher, but the PCC of LAMP1 with OVA was lower in ADJ-treated DCs (**Fig. S2**). This indicates that a significantly higher proportion of EEA1 but not LAMP1 colocalized with antigen in ADJ-treated cells. Collectively, these findings suggest that ADJ preferentially promotes antigen localization to the more alkaline early endosomal compartment.

### ADJ enhances antigen cross-presentation by inducing ROS production and modulating the pH of the antigen-containing compartment by NOX2-dependent mechanisms

Augmented ROS production reduces endosomal cellular acidity and delays antigen degradation, which can in turn enhance antigen cross presentation (Savina et al., 2006). To this end, we used CellROX reagent and the ROS-Glo™ H_2_O_2_ assay to quantify endosomal ROS and H_2_O_2_ production, respectively. Here, we found that total cellular ROS and H_2_O_2_ levels rapidly increased within 1 hour after treatment with ADJ (**Fig. 2C** and S3). Next, we asked whether cellular ROS regulated intracellular acidity using Lysotracker, which has been used for visualizing acidic compartments. Within 1 hour of ADJ treatment, there was a substantive reduction of MFI for Lysotracker, indicating that ADJ reduces intracellular acidity in DCs (**Fig. 2D**). These data collectively suggest that ADJ reduces intracellular acidity by promoting ROS production in DCs.

In order to test the importance of ADJ-induced ROS in cross-presentation, we employed a combination of pharmacological and genetic approaches. First, pharmacological inhibition of NADPH-oxidase complex2 (NOX2) assembly in ADJ-treated DCs using apocynin markedly reduced cross-presentation-dependent stimulation of B3Z cells (**Fig. 2E**). Unlike strong B3Z activation by ADJ-treated DCs from WT mice, ADJ could not augment cross-presentation by DCs deficient for the ROS-inducing NOX2 complex component gp91 (**Fig. 2E**). These data support the idea that maintenance of alkaline environment in the endosomes established by NOX2-driven ROS might be critical for ADJ-mediated cross-presentation.

### ADJ-induced cross-presentation requires proteasomal processing and TAP1 transporters

MHC-I binding peptides can be generated either by phagosomal residential cathepsins, or by cytosolic proteasomes. In order to dissect the pathways required for cross-presentation of OVA-derived peptides by ADJ-treated DCs, we inhibited proteasomal or lysosomal activities using epoxomycin (proteasomal inhibitor) or leupeptin (general cathepsin inhibitor), respectively. Here, ADJ-driven cross-presentation was effectively abrogated by epoxomycin, while it was augmented by leupeptin, as compared to vehicle controls (**Fig. 2F**). This data suggested that ADJ-driven antigen cross-presentation requires proteasomes, not lysosomes. Next, we examined whether ADJ can enhance proteasomal activities in DCs; we found that the activities of 20S proteasome subunits were not affected by ADJ treatment (**Fig. S4**). Thus, ADJ-mediated cross-presentation requires cytosolic proteasomes, but ADJ does not enhance the proteolytic activities of proteasomes in DCs.

Peptides generated by cytosolic proteasomes require TAP transporters to access MHC I molecules in ER or ER-Golgi intermediate compartment (ERGIC) (Cebrian et al., 2011, Van Kaer et al., 1992). We tested the extent to which ADJ-induced cross-presentation is dependent upon TAP transporters in DCs. ADJ-mediated cross-presentation of OVA peptides and antigen recognition by B3Z cells were completely abolished in the absence of TAP1 in DCs (**Fig. 2G**), suggesting that ADJ-mediated cross-presentation requires TAP1 for loading peptides on to MHC I. Collectively, our data suggest that ADJ-induced cross-presentation requires endosomes-to-cytosol-proteasomal pathway of cross-presentation.

### ADJ-mediated cross-presentation requires endosomal antigen leakage mediated by lipid peroxidation

The cytosolic pathway of cross-presentation requires internalized antigens to escape from endosome into cytosol, which are subsequently degraded by cytosolic proteasomes (Kovacsovics-Bankowski and Rock, 1995). Since we observed that ADJ-induced cross-presentation requires proteasomes as antigen processing organelles (**Fig. 2F**), we questioned whether ADJ promotes antigen translocation from endosomes into cytosol. Endosomal leakage in DCs was visualized by measuring cellular apoptosis, resulting from release of horse cytochrome C (cytc) into the cytosol (Lin et al., 2008). After 24 hours of treatment of DCs with cytc in the presence or absence of ADJ, we found that the percentages of annexin-V positive cells were 4 times higher among cells exposed to ADJ+cytc, compared to cytc or ADJ only treated DCs. These data suggest that ADJ likely induced cytc escape from endosomes into cytosol, resulting in DC apoptosis (**Fig. 2H**).

Recently, it was reported that NOX2-driven ROS can cause endosomal lipid peroxidation and release of antigen from leaky endosomes into the cytosol of DCs (Dingjan et al., 2016). Stemming from our results of endosomal leakage in ADJ-treated DCs (**Fig. 2H**), we examined whether ADJ causes lipid peroxidation, which in turn promoted antigen leakage from endosomes. First, we quantified lipid peroxidation in ADJ-treated DCs using a radiometric dye, BODIPY 581/591 C11, which displays a shift in peak fluorescence emission from red to green upon oxidation by lipid hydroperoxides. Within 1 hour after ADJ treatment, there was a marked shift of BODIPY 581/591 C11 from red to green (**Fig. 2I**). Moreover, pharmacological inhibition of lipid peroxidation using α-tocopherol, a lipid-soluble antioxidant which selectively prevents lipid peroxidation by scavenging free electrons (Girotti, 1998), significantly reduced ADJ-driven cross-presentation and activation of B3Z cells (**Fig. 2I**). Hence, our data illustrate that ADJ-mediated cross-presentation requires ROS, and likely lipid peroxidation for effecting endosomal antigen leakage.

### ADJ enhances cross-presentation without glycolytic reprogramming of DCs via the Akt-mTORC1-KLF2-HIF-1α axis

Typically, catabolic metabolism in resting DCs is characterized by oxidative phosphorylation (OXPHOS) fueled by fatty acid oxidation (FAO) and limited glycolysis (Krawczyk et al., 2010, Gotoh et al., 2018, Zuo and Wan, 2019). During early DC activation by TLR agonists, DCs augment both aerobic glycolysis and OXPHOS to support the anabolic demands required for expansion of the ER and Golgi apparatus, *de novo* fatty acid (FA) synthesis, and production of inflammatory cytokines. This early glycolytic reprogramming by TLRs is required for upregulation of co-stimulatory molecules, production of pro-inflammatory cytokines, CCR7 oligomerization, and priming T cells (Everts et al., 2014, Guak et al., 2018). Subsequently, DCs inhibit OXPHOS via NO and rely on aerobic glycolysis for their survival, especially after sustained exposure to TLR-agonists (Everts et al., 2012, Blanco-Perez et al., 2019). Based on the augmented ability of ADJ-treated DCs to activate T cells, we hypothesized that metabolic reprogramming plays a distinctive role in this process.

First, we asked whether ADJ engages aerobic glycolysis, as a key source of carbon for metabolic functions that enhance antigen presentation by DCs via the Akt-mTORC1-HIF-1α signaling axis. As shown in **Fig. 3A**, LPS stimulation potently triggered phosphorylation of p70S6K and Akt within 60 minutes, but treatment with ADJ failed to do the same. Prolonged glycolytic reprogramming involves the induction and stabilization of hypoxia-inducible factor (HIF-1α), that in turn leads to the production of nitric oxide and suppression of OXPHOS (Jantsch et al., 2008, Wilson et al., 2014). Using DCs from HIF-1α luciferase reporter mice (Safran et al., 2006), we compared ADJ and LPS for HIF-1α induction. Data in **Fig. 3B** show that only stimulation with LPS, but not ADJ, induced HIF-1α in DCs. Kruppel-like factor 2 (KLF2) is a transcription factor that inhibits the expression and transcriptional activity of HIF-1α, and downregulation of KLF2 is associated with engagement of glycolysis in immune cells (Mahabeleshwar et al., 2011). As another measure of glycolytic reprogramming, we assessed KLF2 expression levels using BMDCs from KLF2-GFP reporter mice (Skon et al., 2013). High levels of KLF2 were detected in unstimulated DCs, and KLF2 expression was significantly downregulated in LPS-stimulated DCs, but not in ADJ-treated DCs (**Fig. 3C**). To confirm these findings *in vivo*, we immunized KLF2-GFP mice with OVA, ADJ+OVA or LPS+OVA and examined KLF2 expression in DCs in draining lymph nodes. As shown in **Fig. 3D**, DCs from mice immunized with LPS+OVA, but not ADJ+OVA, showed significant downregulation of KLF2 expression, as compared to DCs from OVA only mice. Thus, unlike LPS, ADJ failed to engage the Akt-mTORC1-KLF2-HIF-1α signaling pathway in DCs.

**Figure 3.**
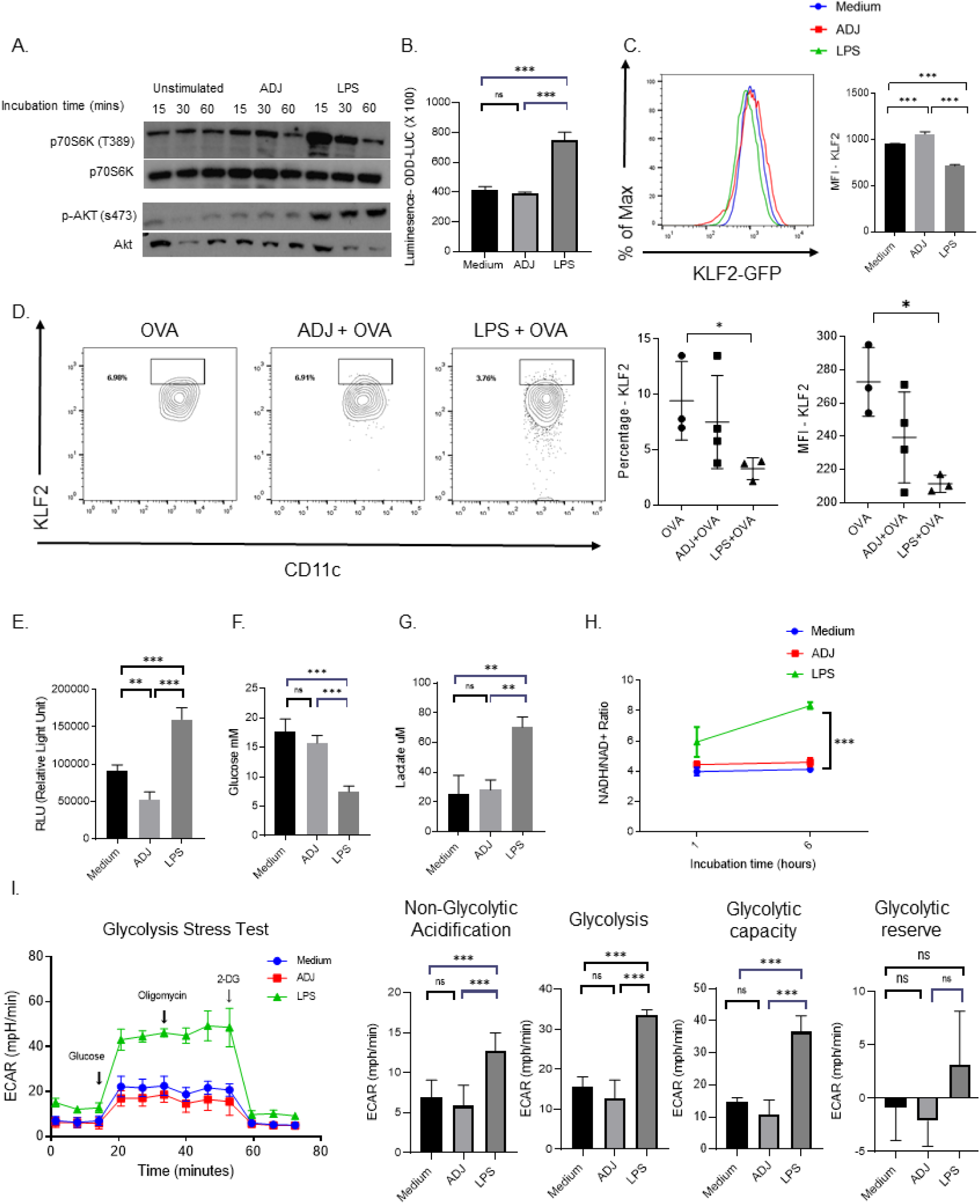
Carbomer-based adjuvant rewires DC metabolism without engaging aerobic glycolysis driven by the Akt-mTORC1-KLF2-HIF1α signaling axis. (A) Immunoblot for total AKT and AKT phosphorylated at Ser473 or total p-S6K and p-S6K phosphorylated at Thr389 in BMDCs following 15, 30, 60 mins of incubation ± ADJ or LPS. (B) HIF1α induction was measured by quantifying luciferase activity in DCs derived from ODD-Luc mice; DCs were stimulated ± ADJ or LPS for 24 h. (C) *In vitro* KLF2 expression in DCs; DCs from KLF2-GFP mice were treated ± ADJ or LPS for 24 hour, and GFP expression was assessed by flow cytometry (D) KLF2-GFP reporter expression (gated on CD11c^HI^/MHC-II^HI^ cDC population) was measured in DCs from DLNs after 24 h of vaccinatio*n in vivo*. (E) Intracellular ATP was quantified in unstimulated, ADJ- or LPS-stimulated DCs at 24 h. (F-G) Extracellular glucose level and secreted lactate in cell culture supernatant was measured in cultures of unstimulated, ADJ-, and LPS-stimulated DCs at 24 h. (H) NADH/NAD+ ratio in resting, ADJ- or LPS-stimulated DCs at the indicated time-point. (I) DCs were treated ± ADJ or LPS for 24 h; real-time ECAR was determined during sequential treatments with glucose, oligomycin, and 2-DG. Quantification of basal glycolysis (ECAR), glycolytic capacity and glycolytic reserve. The glycolytic reserve capacity of cells is the difference between ECAR before and after addition of oligomycin. Data are representative of ≥2 independent experiments. Error bars show SEM; \**P*<0.01; \*\**P*<0.001; \*\*\**P*<0.0001 (Student’s t-test and one-way ANOVA).

To directly investigate the effect of ADJ on DCs’ glycolytic metabolism, we quantified intracellular ATP levels in DCs treated with ADJ or LPS. LPS-treated DCs, but not ADJ-treated DCs contained higher levels of ATP than untreated DCs (**Fig. 3E**). As a measure of glycolysis in ADJ- and LPS-treated DCs, we quantified glucose consumption and lactate production *in vitro*. The extracellular concentrations of glucose and lactate were unaffected by in ADJ stimulation, suggesting that ADJ did not alter glucose utilization or lactate production by DCs (**Fig. 3F and 4G**). Further, we did not find significant alteration of the NADH/NAD+ ratio in ADJ-treated DCs, which suggests that ADJ exposure did not cause a cellular redox imbalance (**Fig. 3H**). Lastly, we quantified the functional glycolytic capacity of ADJ-stimulated DCs using the glycolysis stress test. We found that neither the glycolytic capacity nor the glycolytic reserves were altered in ADJ-treated DCs, as compared to those in unstimulated DCs, while LPS up-regulated both glycolytic capacities and reserves in DCs (**Fig. 3I**). Together, data in **Fig. 3** strongly suggest that ADJ-mediated cross-presentation occurs independent of enhanced aerobic glycolysis.

**Figure 4.**
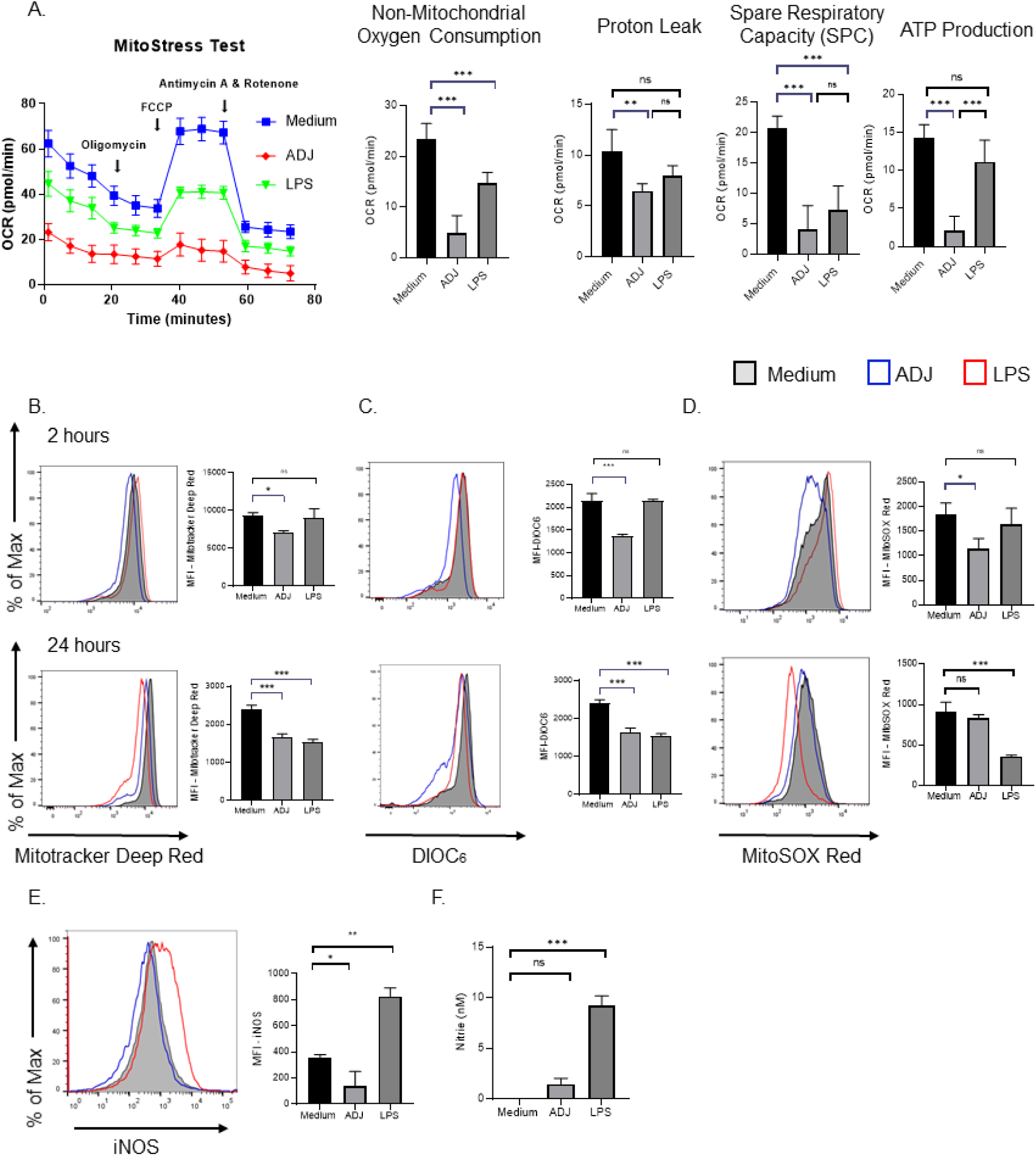
Carbomer-based adjuvant suppresses oxidative phosphorylation in DCs by iNOS-independent mechanisms. (A) BMDCs were treated ± ADJ or LPS for 24 h; real-time OCR was determined during sequential treatments with oligomycin, FCCP, and antimycin-A/rotenone. Quantification of basal respiration, maximal respiration and spare respiratory capacity (SRC). SRC was calculated as the difference in OCR after addition of FCCP (2) and OCR before the addition of oligomycin (B-D) BMDCs were treated ± ADJ or LPS for the indicated time periods (2 or 24 h) and stained with Mitotracker, DiOC_6_, MitoSOX to quantify mitochondrial mass, membrane potential or mitochondrial superoxide, respectively. (E) Culture supernatants from DCs ± ADJ or LPS were analyzed for nitrite levels. (F) iNOS expression in unstimulated, ADJ or LPS-stimulated DCs at 24 h, was determined by intracellular staining. Data are representative of ≥2 independent experiments. Error bars are SEM; \**P*<0.01; \*\**P*<0.001; \*\*\**P*<0.0001 (Student’s t-test and one-way ANOVA).

### ADJ disengages OXPHOS in DCs, independent of iNOS induction

Because ADJ failed to augment glycolysis (**Fig. 3**), we next interrogated whether ADJ engaged OXPHOS as an alternative metabolic pathway. By performing extracellular flux analysis, we measured alterations in oxygen consumption in real time (**Fig. 4A**). At 24 hours after stimulation, the mitochondrial oxygen consumption rate (OCR) was highest for unstimulated DCs (Watson et al., 2019) and mitochondrial OCR was lower for LPS-treated DCs. Notably, baseline OCR for ADJ-treated DCs was markedly lower and failed to show detectable increase, following addition of FCCP, a potent mitochondrial un-coupler that disrupts ATP synthesis (**Fig. 4A**). To further assess ADJ’s effects on mitochondrial metabolism, we measured mitochondrial content, membrane potential (Ψm) and mitochondrial superoxide production (mROS). Within 2 hours of ADJ treatment, we observed a drastic reduction in mitochondrial content, Ψm, and mROS, in comparison to both resting and LPS-stimulated DCs (**Fig. 4B-D**). Interestingly, loss of mitochondrial functions persisted over 24 hours in ADJ-treated cells, indicating that ADJ suppressed mitochondrial functions at both early and late stages of stimulation. These data are consistent with a decrease in spare respiratory capacity (**Fig. 4A**) in ADJ-stimulated DCs, and support the notion that ADJ-mediated metabolic programming includes a profound decline in mitochondrial activity.

One of the mechanisms for inhibiting mitochondrial functions in DCs is the induction of nitric oxide (NO), which interferes with electron transport chain by blocking oxygen consumption and ATP production (Amiel et al., 2014). We probed whether impairment of mitochondrial function by ADJ was linked to NO induction in DCs. The cellular levels of inducible nitric oxide (iNOS) (**Fig. 4E**) and extracellular nitrite levels (**Fig. 4F**) in the supernatant of ADJ-treated DCs did not vary, in comparison to unstimulated DCs, while LPS-stimulated DCs contained elevated levels of cellular iNOS and extracellular nitrite. In summary, these data collectively suggest that ADJ disengages mitochondrial functions by mechanisms independent of iNOS.

### ADJ promotes intracellular lipid body formation in DCs, which is required for ADJ-mediated cross-presentation

Next, we explored how ADJ might enhance cross-presentation independent of aerobic glycolysis in DCs. The formation of intracellular LBs was shown to be critical for efficient cross-presentation in DCs (Bougneres et al., 2009, den Brok et al., 2016). Therefore, we determined whether LBs are also essential for ADJ-mediated cross-presentation. To assess as to whether ADJ can induce intracellular formation of LBs in DCs, we stained unstimulated, LPS-, or ADJ-treated DCs with BODIPY 493/503, which stains neutral lipids. Neutral lipids were barely detected in resting DCs, but ADJ or LPS-stimulated DCs contained abundant levels of neutral lipids, as indicated by an increase in MFI of BODIPY 493/503 in a flow cytometer or as visualized by confocal microscopy (**Fig. 5A-B**). To understand mechanisms underlying the increased intracellular content of neutral lipids in DCs, we examined whether ADJ altered FA uptake by promoting expression of the scavenger receptor CD36. Intriguingly, ADJ increased the expression of CD36 in ADJ-stimulated DCs, but LPS downregulated CD36 expression in DCs (**Fig. 5C**).

**Figure 5.**
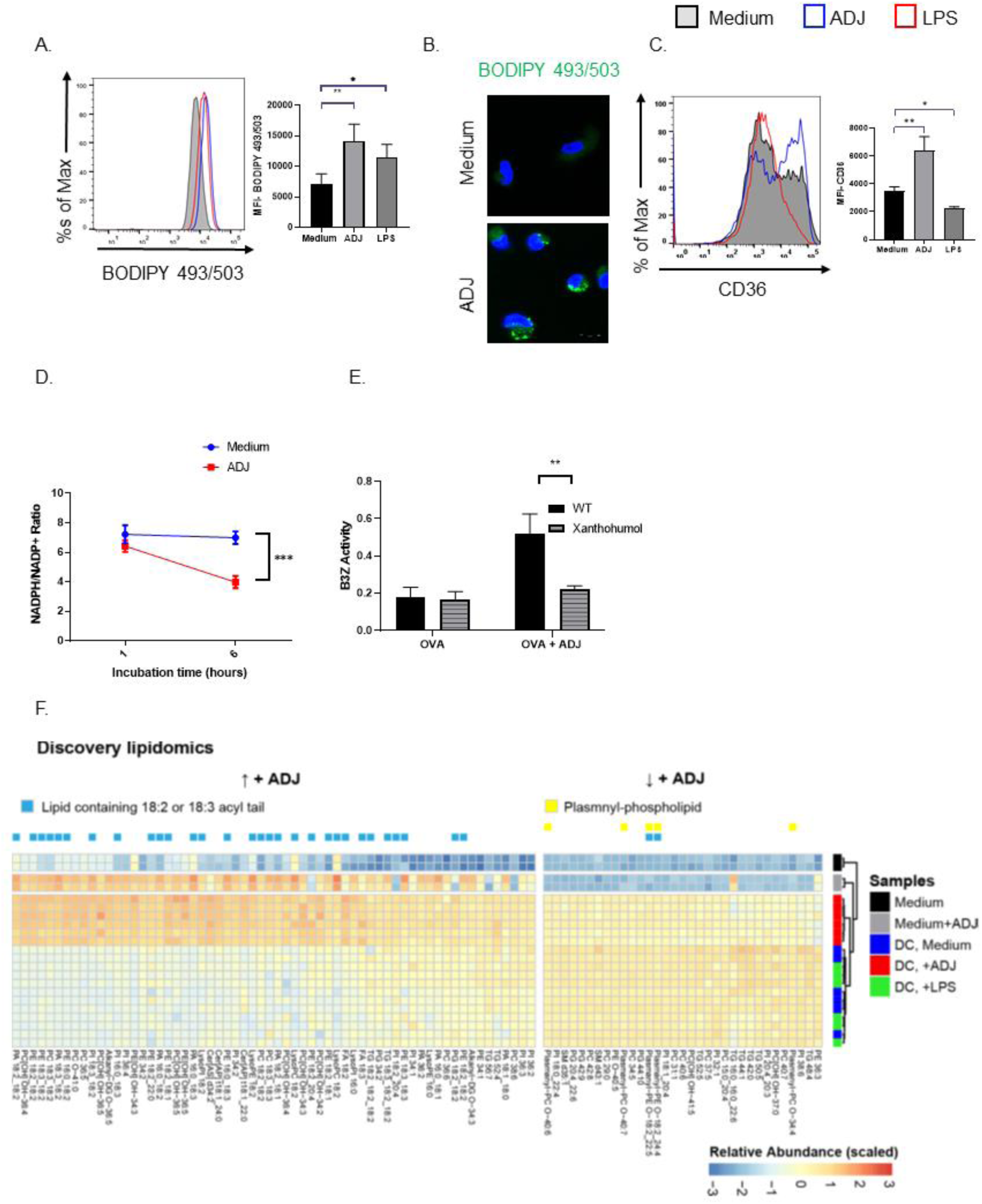
Carbomer-based adjuvant induces intracellular lipid body formation and alters intracellular lipidomes in DCs. (A) Quantification of neutral lipid droplets by flow cytometry and presented as the geometric MFI. (B) Confocal microscopy of DCs treated with ADJ for 24 h, stained with BODIPY, and counter-stained with DAPI. Scale bars, 10 μm. (C) FACS analysis of CD36 expression in unstimulated, and ADJ- or LPS-stimulated DCs after 24 h. (D) NADPH/NADH+ ratio in resting, ADJ- or LPS-stimulated DCs. (E) β-gal production by B3Z cells after co-incubation with ADJ/OVA-treated BMDCs pre-treated with OVA ± ADJ in the presence/absence of xanthohumol for 5 h. (F) Heat-map of lipids from media or DCs stimulated with ADJ ± LPS; lipids were filtered for *P* < 0.05 (ANOVA for +ADJ treatment, Tukey post-hoc corrected p-values) and abundance was scaled across samples. Lipids that increased with +ADJ were enriched with 18:2 or 18:3 acyl-chains (blue bar). Lipids that were decreased with +ADJ contained more plasmanyl-phospholipids (yellow bar). Abbreviations: TG, triglyceride; DG, diglyceride; FA, fatty acid; PA, phosphatidic acid; PC, phosphatidylcholine; PE, phosphatidylethanolamine; PG, phosphatidylglycerol; PI, phosphatidylinositol; SM, sphingomyelin; Cer [AP], alpha-OH fatty acids with pytosphingosine; Cer [AS], alpha-OH fatty acids with sphingosine. Data are representative of ≥2 independent experiments Error bars show SEM; \**P*<0.01; \*\**P*<0.001; \*\*\**P*<0.0001 (Student’s t-test and one-way ANOVA).

It has been reported that glucose-derived pentose phosphate pathway (PPP), an offshoot of the glycolytic pathway, is critical for generation of LBs and pro-inflammatory cytokines upon TLR-stimulation in DCs (Everts et al., 2014). We did not observe increased glycolytic flux in ADJ-stimulated cells to fuel PPP (**Fig. 5I**), but we hypothesized that the NADPH/NADP+ ratio will be altered in ADJ-treated cells because imported FAs need to be activated before incorporation into triglycerides by esterification with coenzyme A, through a reaction catalyzed via fatty acyl-CoA synthetase. As an indirect measure of NADPH/NADP+ ratio, we quantified optical redox ratio (NAD(P)H/NAD(P)H + FAD ratio) in ADJ-stimulated cells using optical multiphoton microscopy (Walsh et al., 2013). We discovered a significant drop in the redox ratio in ADJ-stimulated cells than in unstimulated cells (**Fig. S5**). To further distinguish NADPH/NAD+ from NADH/NAD+ ratio, we used a bioluminescence-based assay and confirmed that ADJ-stimulated cells displayed lower NADPH/NADP+ ratio (**Fig. 5D**). Together, these data suggest that ADJ stimulation likely induces LB formation by utilizing intracellular NADPH and imported FAs, independent of glucose-derived *de novo* FAs.

Lastly, to examine whether LB formation is required for ADJ-driven cross-presentation, we treated DCs with xanthohumol, a DGAT 1/2 inhibitor. Notably, treatment of ADJ-treated DCs with xanthohumol significantly inhibited cross-presentation and activation of B3Z cells by ADJ-treated DCs (**Fig. 5E**). Together, this suggests that LBs might be crucial for ADJ-induced DC cross-presentation.

### ADJ alters intracellular lipidomes in DCs and increases accumulation of 18:2 and 18:3-containing lipids

The saturation and oxidation status of lipid might modulate cross-presentation by DCs (Veglia et al., 2017, Herber et al., 2010, Ramakrishnan et al., 2014, Lee et al., 2004). However, how lipid composition governs DC cross-presentation remains unknown. In order to map changes in lipid species in ADJ-treated cells, we employed a discovery lipidomics mass spectrometry approach. With this approach, we identified 446 unique lipid species across 25 lipid classes. As compared to resting or LPS-treated DCs, ADJ-treated DCs displayed significant increases in lipids containing acyl-chains with linoleic (18:2) or alpha-linoleic (18:3) acid (**Fig. 5F**), which appear to be constituents of the adjuvant ADJ (**Fig. 5F**, gray bars); media with ADJ also contained an increased abundance of 18:2 and 18:3-containing lipids, as compared to media alone. In addition, ADJ treatment also led to increases in ceramides containing alpha-hydroxy fatty acids (Cer[AP] and Cer[AS]) and decreased abundance of plasmanyl-phospholipids. Hence, our lipidomic profiles further highlight the global lipid changes induced by ADJ treatment, characterized by increased abundance of linoleic and alpha-linoleic acyl-chains within phospholipids and triglycerides, likely resulting in changes in membrane fluidity. Together, our data suggest that changes in lipid composition could be critical for ADJ-mediated cross-presentation in DCs.

### ADJ elicits T cell-based protective immunity against viral and intracellular bacterial infections in vivo

Lastly, we examined the adjuvanticity of ADJ as a T cell-based vaccine by questioning whether ADJ augments activation and expansion of antigen-specific CD8 T cells *in vivo* following vaccination with the experimental subunit antigen chicken ovalbumin (OVA). As shown in **Fig. 6A**, formulation of OVA with ADJ potently augmented the percentages of activated OVA-specific CD8^+^ T cells in spleen, while OVA alone displayed poor immunogenicity. Next, we evaluated the ability of ADJ-based vaccine to confer T cell-based protection to pathogens *Listeria monocytogenes (LM)* or vaccinia virus (VV) in mice (Roberts et al., 1993, Ladel et al., 1994, Harty et al., 2000). 40 days after last vaccination, mice were challenged with either recombinant LM-expressing OVA (LM-OVA) or recombinant VV-expressing OVA (VV-OVA) (McCabe et al., 1995, Shen et al., 1998). After LM-OVA or VV-OVA challenge, we enumerated recall OVA-specific CD8 T cell responses in spleen or in lung, and LM-OVA or VV-OVA burden in various tissues. After challenge, higher numbers of OVA SIINFEKL epitope-specific CD8 T cells were detected in spleens or lungs of ADJ+OVA-vaccinated mice, as compared to those in unvaccinated mice (**Fig. 6B-6C**). Consistent with potent antigen-specific recall CD8 T cell responses in ADJ+OVA mice, LM-OVA and VV-OVA burdens in tissues of ADJ+OVA group were markedly lower than in unvaccinated controls (**Fig. 6D-E**). Together, data in **Figure 6** clearly demonstrated that ADJ-based subunit vaccine provided protective T-cell-based protective immunity against bacterial and viral pathogens by promoting cross-presentation of antigen to SIINFEKL-specific CD8 T cells *in vivo*.

**Figure 6:**
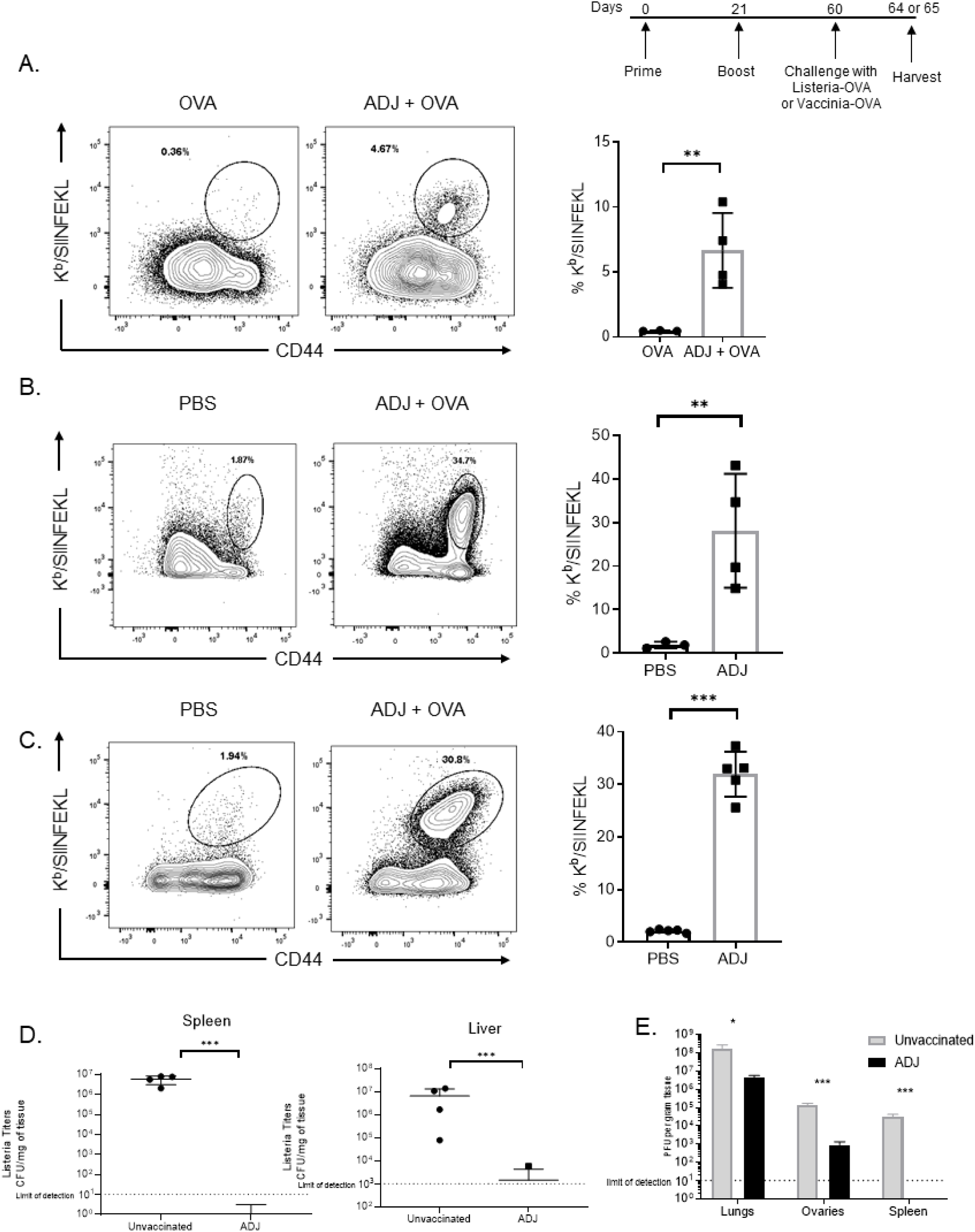
Carbomer adjuvant-based subunit vaccine induces potent CD8 T cell responses and protects against Listeria and vaccinia challenge *in vivo*. (A) C57BL/6 mice were vaccinated by SQ injection of 10ug OVA ± 5% ADJ, and boosted 21 days later. At day 8 after vaccination, activated OVA SIINFEKL-specific CD8 T cells) were quantified in spleens by flow cytometry; plots are gated on total CD8 T cells. (B-D) Forty days after vaccination, mice were challenged with 1.7×10^5^ CFUs of virulent LM-OVA intravenously or 2×10^6^ PFUs of VV-OVA intranasally; unvaccinated mice (PBS) were challenged as controls. (B) 4 days after LM-OVA challenge, activated OVA SIINFEKL-specific CD8 T cells (gated on CD8 T cells) were quantified in spleen by flow cytometry. (C) 5 days after VV-OVA challenge, activated OVA SIINFEKL-specific CD8 T cells (gated on CD8 T cells) were quantified in lung by flow cytometry. (D-E) At day 4 (for LM-OVA) or 5 (for VV-OVA) after challenge, LM-OVA burden (spleen and liver) or VV-OVA titers (lungs, ovaries, and spleens) were quantified. Data are representative of ≥2 independent experiments Error bars are SEM; \**P*<0.01; \*\**P*<0.001; \*\*\**P*<0.0001 (Student’s t-test and one-way ANOVA).

## DISCUSSION

In this manuscript, we report that a carbomer-based nano-emulsion adjuvant, ADJ, promoted memory T cell-dependent protective immunity against intracellular pathogens, *Listeria monocytogenes* and vaccinia virus. Consistent with strong induction of CD8 T cell immunity to subunit antigens *in vivo*, ADJ promoted DC cross-presentation *in vitro*. Mechanistic investigations of antigen processing and the metabolic basis for adjuvant action demonstrate that ADJ promoted multiple aspects of antigen cross-presentation in DCs, in the apparent absence of metabolic switch from catabolic to anabolic phenotype. This constitutes a novel mechanism because the current axiom suggests that engagement of active aerobic glycolysis in DCs is necessary for their activation and ability to stimulate CD8 T cells (Pearce and Everts, 2015).

How does ADJ enhance DC cross-presentation and potentiate CD8 T cell immunity *in vivo*? Effective cross-presentation requires antigen targeting to alkaline intracellular compartment, slow degradation of antigens by proteases, and translocation of endosomal antigens into cytosol. Interestingly, we discovered that ADJ-treated DCs contained greater amounts of degraded antigens, compared to untreated DCs, but this was not linked to elevated antigen uptake. Increased antigen cross presentation here seems to be the result of greater induction of ROS by ADJ that presumably leads to alkalization, attenuated antigen degradation, and accumulation of partially degraded antigens in alkaline endosomes. Thus, one mechanism by which ADJ might enhance cross-presentation is by delaying antigen degradation and promoting partially degraded antigen accumulation in DCs, which in turn sustains antigen presentation by DCs.

Our studies also demonstrate that ADJ-treated DCs utilize an endosomal-to-cytosol pathway of cross-presentation, as indicated by augmented endosomal protein leakage into cytosol and abolishment of ADJ-driven cross-presentation by pharmacological inhibition of cytosolic proteasomes. The molecular mechanisms underlying the translocation of endosomal antigens into cytosol during cross-presentation remains controversial. One possible mechanism is explained by the transporter hypothesis, in which exogenous antigens are unfolded by gamma-interferon-inducible lysosomal thiol reductase (GILT) in the phagosome and transported into cytoplasm by ER-associated degradation (ERAD) machinery or chaperone-mediated transport by Hsp90 (Kato et al., 2012, Singh and Cresswell, 2010, Imai et al., 2005). Another potential mechanism is endosomal membrane disruption, induced by NOX2-dependent ROS production that serves as precursors for lipid peroxides in the endosomes. In our studies, we found that: (1) ADJ strongly induced ROS production; (2) NOX2 deficiency and pharmacological inhibition of NOX2 assembly abolished cross-presentation; (3) antioxidant tocopherol diminished ADJ-driven cross-presentation. Therefore, our data favor the endosomal disruption model, where ADJ-induced ROS might have a dual function in cross-presentation: increase endosomal pH to delay antigen degradation and promote antigen translocation into cytosol by lipid peroxidation.

Only recently, it was discovered that TLR-driven metabolic shift to anabolic metabolism is an integral component of the activation program that is required to activate naïve T cells (Pearce and Everts, 2015). While the metabolic basis of how TLR agonists support DC effector functions is well characterized, the exact metabolic roles of many immune adjuvants in dictating DC activation and antigen presentation remain poorly understood. Our data suggest that the two important energy yielding metabolic pathways, glycolysis and OXPHOS, were minimally engaged or inactive in ADJ-treated DCs, which is indicative of a unique cellular state of metabolic quiescence during cross-presentation. How ADJ-treated DCs effectively stimulate T cells *in vivo* without the need to switch to glycolysis or to trigger enhanced mitochondrial metabolism remains unclear. Recently, it was reported that DCs contain intrinsic glycogen, which can be readily catabolized into glucose upon LPS stimulation to fuel intracellular glucose (Thwe et al., 2017). Unlike FAO, which requires a substantial number of functional mitochondria, glycogenolysis occurs in the cytoplasm. Because ADJ treatment results in a loss of mitochondrial functions and minimal engagement of glycolysis in DCs, it is plausible that ADJ-treated DCs can catabolize intrinsic glycogen into glucose to sustain their survival in the absence of functional mitochondria. Follow-up studies should evaluate whether ADJ-treated DCs utilize intracellular glycogen reserves during metabolic quiescence and/or whether glycogenolysis is required for ADJ-mediated DC cross-presentation.

Despite a ‘hypometabolic’ state, ADJ-treated DCs displayed effective antigen cross-presentation and stimulation of CD8 T cell responses *in vivo*. However, this leads to the question of why ADJ-stimulated DCs adapt to this particular type of metabolism that is metabolically inefficient, at least in terms of ATP production. A recent study suggests that high levels of glucose represses DC-induced T cell responses by engaging the mTORC1-HIF1α/iNOS pathway in DCs (Lawless et al., 2017). In their studies, limiting glucose availability to DCs enhanced T cell responses by diminishing competition for glucose from DCs in the immune microenvironment. It is plausible that ADJ drives DCs to a metabolic state that is less dependent upon glucose-driven catabolic pathways, which enables DC outputs tailored for effective cross-presentation to T cells. For instance, conventional processing of exogenous antigens involves acidification of the lysosomal compartment, which is required for efficient generation of peptides for loading into MHC II and that is dependent on activity of ATP-driven proton pumps (Trombetta et al., 2003).

During this process, lysosomal V-ATPase complex is assembled by intracellular H^+^ ions generated by aerobic glycolysis (resulting from oxidation of NADH into NAD^+^ and H^+^ ions), leading to an increase in lysosomal acidity. Notably, ADJ did not trigger redox imbalance of NADH/NAD+ ratio in DCs, but only reduced intracellular ATP production. Thus, it is plausible that maintenance of a low metabolic state by reducing a key carbon source for generating ATP and keeping balanced redox ratio might be essential for reducing intracellular acidity by inhibiting lysosomal V-ATPase assembly in DCs.

The divergent functions of inflammasome activation in shaping adaptive immunity have been demonstrated for other vaccine adjuvants, such as Alum and ISCOM (Wilson et al., 2014). Similar to these adjuvants, ADJ induced inflammasome activation in DCs. This finding was unexpected, especially because LPS-induced succinate stabilized HIF-1α in macrophages, which is known to be required for inflammasome activation under inflammatory conditions (Tannahill et al., 2013). However, ADJ did not promote stabilized-HIF1α accumulation under normoxia in DCs. It has been also documented that phagolysosomal destabilization after adjuvant phagocytosis, such as Alum and Carbopol, is an important step in inflammasome activation (Gartlan et al., 2016, Hornung et al., 2008). While we have not directly characterized the intracellular location of ADJ in DCs, ADJ increased lysosomal pH, which in turn may result in lysosomal stabilization. How ADJ drives inflammasome activation independent of lysosomal rupture or HIF-1α-stabilization is unknown, but inflammasome activation has been closely associated with enhanced cross-presentation in multiple studies (Sokolovska et al., 2013, Li et al., 2019). Despite strong evidence of inflammasome activation by ADJ, caspase 1 deficiency did not significantly affect ADJ’s ability to stimulate antigen-specific CD8 T cell responses *in vivo*, indicating that ADJ-mediated cross-presentation occurs independent of inflammasome activation. Such finding is consistent with a previous report, in which induction of OVA-specific antibody titers by Carbopol, a polyanionic carbomer, was not significantly affected in NLRP3 or caspase 1-deficient mice (Gartlan et al., 2016). Future studies are warranted to determine how ADJ triggers inflammasome activation in DCs and the role of inflammasome activation in engendering protective cell-mediated immunity *in vivo*.

The causative link between the formation of intracellular LBs and cross-presentation efficiency by IFN-γ and ISCOMs has been recently established (Bougneres et al., 2009, den Brok et al., 2016). Consistent with previous findings, our studies also suggest that ADJ-induced cross-presentation requires intracellular LB formation in DCs. However, how LBs exert their effects on DC cross-presentation remains undetermined. While we have not yet examined whether ADJ-aided LBs directly affect antigen processing, our lipidomic studies highlight the unique composition of lipids within ADJ-treated DCs, which shows accumulations of 18:2 and 18:3 acyl tails and ceramides, presumably due to uptake of ADJ itself. Because ADJ upregulates CD36 expression in DCs, it is conceivable that ADJ enhances the uptake of external lipid without *de novo* fatty acid synthesis derived from glucose (Rosas-Ballina et al., 2020). In line with low metabolic profiles in ADJ-stimulated DCs, the formation of intracellular LBs using external fatty acids could be a more efficient pathway to store intracellular lipids since it requires minimal energy and overall metabolic activities. The global intracellular lipidome modified by ADJ suggests a possible explanation for ADJ-induced cross-presentation, presumably by regulating antigen export to cytosol during cross-presentation. For example, an increase in double-bonds within phospholipids could directly increase membrane fluidity, leading to an increase in endosomal antigen leakage (Shaikh and Edidin, 2008, Shaikh and Edidin, 2006). An enrichment in certain ceramides could also contribute to the formation of lipid rafts, which are known to be critical for regulation of endosomal NOX2 assembly (Rao Malla et al., 2010). Thus, enrichment of certain lipid classes in the endosomes mediated by ADJ, such as unsaturated phospholipids and ceramides, could be another important rate-liming step for antigen export to the cytosol, in addition to ROS-driven endosomal disruption.

In summary, we have identified a carbomer-based adjuvant (ADJ) that elicits protective CD8 T cell responses to soluble subunit antigens, protects against viruses and an intracellular bacterial pathogen, and enhances cross-presentation by the cytosolic pathway. The most striking finding was that ADJ-stimulated DCs, which are highly efficient in cross-presenting antigens, exhibited a distinct metabolic state that is characterized by minimum glycolytic activity, low mitochondrial respiration, and intracellular LB formation. Thus, our model challenges the prevailing metabolic paradigm by suggesting that retaining DCs in a quiescent state is a unique mechanism to regulate efficient DC cross-presentation. Our findings have significant implications in understanding the mechanism of action of adjuvants and development of safe and effective vaccines that elicit potent T cell-based immunity against infectious diseases, such as HIV, TB and malaria.

## STAR METHODS

### KEY RESOURCES TABLE

| REAGENT or RESOURCE | SOURCE | IDENTIFIER |
| --- | --- | --- |
| Antibodies |  |  |
| Hamster anti-CD11c-BV421-conjugated (N418) | BD Pharmingen | 565452 |
| Hamster anti-CD11c/PE-Cy7-conjugated (N418) | BD Pharmingen | 558079 |
| Rat anti-CD40-BV421-conjugated (3/23) | BD Pharmingen | 562846 |
| Hamster anti-CD80-APC-conjugated (16-10A1) | BD Pharmingen | 560016 |
| Mouse anti-H-2Kb-FITC-conjugated (AF6-88.5) | BD Pharmingen | 562002 |
| Mouse anti-I-A[b]-PE-CF594-conjugated (AF6-120.1) | BD Pharmingen | 562824 |
| Annexin V-PE-conjugated | BD Pharmingen | 556421 |
| Rat Anti-CD4-BUV496-conjugated (GK1.5) | BD Pharmingen | 564667 |
| Rat Anti-CD8-BUV395-conjugated (53-6.7) | BD Pharmingen | 563786 |
| Rat Anti-CD44-BV510-conjugated (IM7) | BD Pharmingen | 563114 |
| Mouse Rat anti-CD86-PE/Cy7-conjugated (GL-1) | Biolegend | 105014 |
| Rat anti-CCR7-PE-conjugated (4B12) | eBioscience | 12-1971-82 |
| Hamster anti-CD36-Alexa Fluor700-conjugated (HM36) | eBioscience | 56-0362-80 |
| APC-conjugated H2-Kb tetramers bearing the ovalbumin peptide SIINFEKL | NIH Tetramer Core Facility at Emory University |  |
| Mouse anti-NOS2 Antibody PE-conjugated (C-11) | Santa Cruz<br>Biotechnology | sc-7271 |
| Rat anti-mouse LAMP1 antibody (1D4B) | Santa Cruz<br>Biotechnology | sc-19992 |
| Purified Anti-Mouse CD16 / CD32 (Fc Shield) (2.4G2) | Tonbo Biosciences | 70-0161-U500 |
| Alexa Fluor 488 goat anti-rabbit IgG (H+L) secondary antibodies | Thermo Fisher Scientific | A11008 |
| Alexa Fluor 568 goat anti-rat IgG (H+L) secondary antibodies | Thermo Fisher Scientific | A11077 |
| Goat anti-Rabbit IgG (H+L) Cross-Adsorbed Secondary Antibody, HRP | Thermo Fisher Scientific | G-21234 |
| Rabbit anti-EEA1 antibody (C45B10) | Cell Signaling<br>Technology | 3288 |
| Rabbit anti-Phospho-Akt (Ser473, Clone: D9E) antibody | Cell Signaling<br>Technology | 4060S |
| Rabbit anti-Akt polyclonal antibody | Cell Signaling<br>Technology | 9272S |
| Rabbit anti-phosphor-p70 S6 kinase (Thr389, Clone: 108D2) | Cell Signaling<br>Technology | 9324S |
| Rabbit polyclonal anti-p70 S6 kinase antibodies | Cell Signaling<br>Technology | 9202S |
| Bacterial and Virus Strains |  |  |
| Recombinant vaccinia virus expressing chicken ovalbumin (VV-OVA) | Provided by Jack Bennink (NIH) | (McCabe et al., 1995) |
| Recombinant *Listeria monocytogenes* expressing chicken ovalbumin (LM-OVA) | Provided by Dr. Hao Shen (University of Pennsylvania) | (Shen et al., 1998) |
| Chemicals, Peptides, and Recombinant Proteins |  |  |
| Adjuplex (endotoxin-free) | Purchased from Empiricon | N/A |
| Hen egg white ovalbumin grade V (OVA) from chicken egg white | MilliporeSigma | A5503 |
| LPS purified from *Escherichia coli* O111:B4 | MilliporeSigma | L2630 |
| Leupeptin | MilliporeSigma | L2884 |
| α-Tocopherol | MilliporeSigma | T3251 |
| Gelatin from cold water fish skin | MilliporeSigma | G7041 |
| Collagenase B | MilliporeSigma | 11088831001 |
| Collagenase D | MilliporeSigma | 11088882001 |
| Cytochrome c from equine heart | MilliporeSigma | C2506 |
| Ovalbumin, Alexa Fluor™ 647 Conjugate | Thermo Fisher Scientific | O34784 |
| DQ™ Ovalbumin | Thermo Fisher Scientific | D12053 |
| CellROX Deep Red reagent | Thermo Fisher Scientific | C10422 |
| Lysotracker Deep Red reagent | Thermo Fisher Scientific | L12492 |
| LiveDead eFlour 780 stain | eBioscience | A10628 |
| BODIPY 493/503 | Thermo Fisher Scientific | D3922 |
| MitoTracker™ Deep Red FM | Thermo Fisher Scientific | M22426 |
| DiOC<sub>6</sub>(3) | Thermo Fisher Scientific | D273 |
| MitoSOX™ Red Mitochondrial Superoxide Indicator | Thermo Fisher Scientific | M36008 |
| Protease Inhibitor Tablets | Thermo Fisher Scientific | PI88666 |
| Pierce™ Phosphatase Inhibitor Mini Tablets | Thermo Fisher Scientific | 88667 |
| Thermo Scientific Halt Protease Inhibitor Cocktails | Thermo Fisher Scientific | 78430 |
| Prolong Gold Antifade | Thermo Fisher Scientific | P36934 |
| DAPI | Thermo Fisher Scientific | D1306 |
| Promega pHAb Amine Reactive Dye | Promega | G9841 |
| Chlorophenol red-β-D-galactopyranoside (CPRG) | Santa Cruz<br>Biotechnology | sc-257242 |
| Apocynin | Santa Cruz<br>Biotechnology | sc-203321 |
| Epoxomycin | Cayman Chemicals | 10007806 |
| Xanthohumol | Cayman Chemicals | 15399 |
| Recombinant murine GM-CSF | Peprtech | 315-03 |
| 16% paraformaldehyde | Electron Microscopy<br>Sciences | 15710-S |
| NP-40 | VWR (Amresco) | M158-50ML |
| TRIS | VWR (Amresco) | 97061-794 |
| Tween-20 | VWR (Amresco) | 97062-332 |
| TRITON™ X-100 | VWR (Amresco) | 97063-864 |
| Glycine | VWR (Amresco) | 0167-1kg |
| Brain Heart Infusion | VWR | 90003-040 |
| Glutaraldehyde | Fisher Scientific | O2957-1 |
| Agar | Fisher Scientific | BP1423-500 |
| Critical Commercial Assays |  |  |
| Cytofix/Cytoperm™ fixation/permeabilization | BD Biosciences | 555028 |
| Glycolysis Stress Kit | Agilent | 103020-100 |
| Mito Stress Kit | Agilent | 103708-100 |
| Glucose-Glo™ Assay | Promega | J6021 |
| Lactate-Glo™ Assay | Promega | J5021 |
| NADH/NAD<sup>+</sup>-Glo™ Assay | Promega | G9071 |
| NADPH/NADP<sup>+</sup>-Glo™ Assay | Promega | G9081 |
| ROS-Glo H<sub>2</sub>O<sub>2</sub> assay | Promega | G8820 |
| Proteasome-Glo™ Chymotrypsin-Like, Trypsin-Like and Caspase-Like Cell-Based Assays | Promega | G1180 |
| CellTiter-Glo® 2.0 Cell Viability Assay | Promega | G9241 |
| Luciferase Assay System | Promega | E4030 |
| Griess' reagent for nitrite quantification | MilliporeSigma | 03553-100ML |
| Bio-Plex Pro Mouse Cytokine 23-plex Assay (Bio-rad) | Bio-Rad | M60009RDPD |
| Image-iT™ Lipid Peroxidation Kit, for live cell analysis | Thermo Fisher Scientific | C10445 |
| Thermo Scientific Pierce BCA Protein Assay | Thermo Fisher Scientific | 23227 |
| Mouse IL-18 ELISA kit | Thermo Fisher Scientific | BMS618-3 |
| Mouse IFN-β ELISA kit | R&D Systems | DY8234-05 |
| Experimental Models: Cell Lines |  |  |
| VERO cells | ATCC | ATCC® CCL-81™ |
| B3Z T cell reporter cell line | Obtained from Dr. Bruce Klein (University of Wisconsin-Madison) | (Karttunen et al., 1992) |
| Experimental Models: Organisms/Strains |  |  |
| C57BL/6J mice | Jackson Labs | # 000664 |
| B6.129S2-Tap1<sup>tm1Arp</sup>/J (TAP1 KO) | Jackson Labs | # 002944 (Van Kaer et al., 1992) |
| B6.129S-Cyb<sup>btm1Din</sup>/J (gp91phox-/NOX2 KO) | Jackson Labs | # 002365 (Pollock et al., 1995) |
| B6N.129S2-Casp1<sup>tm1Flv</sup>/J (Caspase 1 KO) | Jackson Labs, obtained from Dr. JD Sauer (University of Wisconsin-Madison) | # 016621 (Kuida et al., 1995) |
| B6. KLF2-GFP reporter mice | Dr. Stephen Jameson (University of Minnesota) | (Skon et al., 2013). |
| Albino B6. C57BL6J ODD-Luc | Dr. Richard Einstein (University of Wisconsin-Madison) | (Safran et al., 2006), |
| B6.129S6-Nlrp3tm1Bhk/J (NLRP3 KO -/-) | Obtained from Dr.<br>Dmitry Shayakhmetov<br>(Emory University);<br>Kindly provided by Drs.<br>Vishva Dixit and Kim<br>Newton (Genentech -<br>Roche, CA) | # 021302 |
| B6. ASC KO -/- | Obtained from Dr.<br>Dmitry Shayakhmetov<br>(Emory University);<br>kindly provided by Drs.<br>Vishva Dixit and Kim<br>Newton (Genentech -<br>Roche, CA) |  |
| B6. NLRP3 / AIM2 KO | Obtained from Dr.<br>Dmitry Shayakhmetov<br>(Emory University);<br>kindly provided by Dr.<br>Thirumala-Devi<br>Kannegati (St. Jude<br>Childrens Hospital, TN). |  |
| Software and Algorithms |  |  |
| FlowJo v10 | TreeStar | N/A |
| Prism V 7.0 | Graphpad Prism | N/A |
| Bioplex Manager 6.1.1 | Bio-Rad | N/A |
| LAS X software | Leica | N/A |
| Wave Software Version 2.4 | Agilent | N/A |
| CellProfiler | CellProfiler | N/A |
| Lipidex | (Hutchins et al., 2018) |  |
| Compound Discoverer 2.1 | Thermo Fisher Scientific | N/A |
| Other |  |  |
| RPMI-1640 Media | Lonza | 12-702F/12 |
| ACK buffer | Lonza | 10-0548E |
| Penicillin/Streptomycin | Lonza | 17-602F |
| L-Glutamine | Lonza | 12-702F/12 |
| IMDM Media | Lonza | 12-726Q |
| DMEM Media (with low glucose) | Lonza | 12-707F |
| RPMI 1640 without L-Gln or Phenol Red | Lonza | 12-918F |
| PBS | Lonza | 17-516F/24 |
| Fetal Bovine Serum (FBS) - premium | Atlanta Biologicals | S11150 |
| Bovine Serum Albumin | Fisher Scientific | BP1600-1 |
| 70 µm Falcon® Cell Strainers, Sterile, White, Corning | Corning | 352350 |
| PBS powder | Bio Basic | PD0100 |
| RIPA buffer | Cell Signaling Technology | 9806S |
| Surepage™, Bis-Tris, 10x8 Cassette, 8 x 8 gel, 10%, 12 wells | Genscript | M00665 |
| iBlot™ 2 Transfer Stacks, PVDF, regular size | Thermo Fisher Scientific | IB24001 |
| Fetal Bovine Serum, dialyzed, US origin | Thermo Fisher Scientific | A3382001 |
| Trypsin-EDTA (0.05%), phenol red | Thermo Fisher Scientific | 25300120 |
| Goat serum | Thermo Fisher Scientific | 16210064 |
| Non-fat dry milk | LabScientific | BM-0841 |
| ECL Prime Western Blotting Detection Reagent | GE Healthcare | RPN2236 |
| Disposable PD-10 Desalting Column, with<br>Sephadex G-25 | GE Healthcare | 17085101 |
| UltraCruz® Autoradiography Film | Santa Cruz<br>Biotechnology | sc-201696 |
| Coverslips 12mm (# 1.5 thickness) | Warner Instruments | 64-0712 |
| GentleMacs C Tubes | Milltenyi Biotech | 130-096-334 |
| Non-sterile CellTrics® filters | Sysmex | 04-0042-2317 |
| Brilliant Staining Buffer | BD Biosciences | 566385 |
| Equipment |  |  |
| SP8 3X STED super-resolution microscope. | Leica |  |
| GentleMACS dissociator | Miltenyi Biotech |  |
| BD LSRFortessa | BD |  |
| iBlot™ 2 Gel Transfer Device | Thermo Fisher Scientific |  |
| Bioplex 200 systems | Bio-rad |  |
| Spectramax i3x multi-mode microplate reader | Molecular Devices |  |
| A custom-built multi-photon microscope | Bruker | (Walsh and Skala,<br>2015) |
| Acquity CSH C18 column | Waters |  |
| Q Exactive HF Orbitrap mass spectrometer | Thermo Fisher Scientific |  |
| Vanquish Binary Pump | Thermo Fisher Scientific |  |

### CONTACT FOR REAGENT AND RESOURCE SHARING

Further information and requests for resources and reagents should be directed to and will be made available upon reasonable request by the Lead Contact, M. Suresh. This study did not generate new unique reagents.

## METHODS

### Experimental animals

7-12-week-old C57BL/6 (B6), Gp91 (NOX2) -/- (Stock number, 002365), and TAP1 -/- (Stock number: 002944) were purchased from Jackson Laboratory or from restricted-access SPF mouse breeding colonies at the University of Wisconsin-Madison Breeding Core Facility. Caspase 1-deficient and ODD-LUC mice backcrossed to Albino C57BL/6 background provided by Drs. J. D. Sauer, and Richard Eisenstein (University of Wisconsin-Madison), respectively. KLF2-GFP reporter mice were provided by Dr. Stephen Jameson (University of Minnesota). NLRP3-KO and ASC-KO mice were kindly provided by Drs. Vishva Dixit and Kim Newton (Genentech-Roche, CA); NLRP3/AIM2-dKO mice were kindly provided by Dr. Thirumala-Devi Kanneganti (St. Jude Childrens Hospital, TN).

### Ethics statement

All animal experiments were performed in accordance with the protocol (Protocol number V5308 and V5564) approved by the University of Wisconsin School of Veterinary Medicine Institutional Animal Care and Use Committee (IACUC). The animal committee mandates that institutions and individuals using animals for research, teaching, and/or testing much acknowledge and accept both legal and ethical responsibility for the animals under their care, as specified in the Animal Welfare Act (AWA) and associated Animal Welfare Regulations (AWRs) and Public Health Service (PHS) Policy.

### Tissue processing and Flow cytometry

Primary monoclonal antibodies for detecting surface markers/tetramers were used for flow cytometry at 1:200 dilution (provided in the resource table) unless stated otherwise. Spleens and lymph nodes were processed into single-cell suspensions by standard procedures. Briefly, for collagenase-based digestion, tissue samples were digested in 2mg/mL of Collagenase (Collagenase B for lungs; Collagenase D for draining lymph nodes) for 15 minutes at 37C. Single-cell suspensions were first stained for viability with LiveDead eFlour 780 stain (eBioscience) and stained with antibodies diluted in Brilliant Stain Buffer (BD Biosciences) or FACS buffer (1% BSA in PBS), or complete RPMI media (10% fetal bovine serum [FBS] in RPMI) for 30-60 minutes. Intracellular staining of iNOS was performed using Cytofix/Cytoperm™ fixation/permeabilization kit (BD) as previously described (Everts et al., 2012). Samples were acquired with a BD LSR Fortessa (BD Biosciences) and resulting data were analyzed with FlowJo software (TreeStar, Ashland, OR).

### BMDC generation and cell culture

Primary cultures of bone marrow-derived DCs were generated as previously described (Na et al., 2016, Jin and Sprent, 2018). Briefly, femur, tibia, and humerus from the mice were flushed using RPMI-1640 medium supplemented with 1% FBS. BMDCs were maintained in RPMI-1640 medium supplemented with 10% fetal bovine serum, 100 U/ml penicillin G, 100 ug/ml streptomycin sulfate, and 10ng/ml GM-CSF (Peprotech) in 150mm petri dishes. Equal parts of additional media with 10ng/ml of GM-CSF were added on day 3. Loosely attached immature DCs were collected at day 6-7 and subsequently used for the experiment. The B3Z hybridoma was a generous gift from Dr. Bruce Klein (University of Wisconsin-Madison). B3Z cells were maintained in Iscove’s Modified Dulbecco’s media supplemented with 10% FBS, 100 U/ml penicillin G and 100 g/ml streptomycin sulfate.

### Vaccination, Viral titers, and enumeration of Listeria Challenge

C57BL/6 Mice were vaccinated by the subcutaneous route at the tail base with 50ul of the vaccine (containing 5% ADJ and 10ug chicken OVA). Mice were boosted after 21 days of the initial vaccination. At > 40 days after vaccination, mice were challenged intranasally with 2 × 10^6^ recombinant vaccinia virus-expressing OVA. Spleens, lungs, and ovaries were used for viral titers. Tissue samples for viral titers were homogenized in 400ul 1mM pH 8.0 TRIS using bead homogenizer. Clarified supernatant was trypsinized in Trypsin-EDTA (0.05%) and titrated in complete RPMI on Vero cells. *Listeria monocytogenes* expressing chicken ovalbumin (LM-OVA) was provided by Dr. Hao Shen (University of Pennsylvania School of Medicine). Mice were infected intravenously with 1.7 x 10^5^ CFUs of LM-OVA. To quantify Listeria burdens, tissues were homogenized in gentleMACS C-Tubes via gentleMACS dissociator. Organs were processed in sterile 0.1% Nonidet-P40 + PBS in gentleMACS C Tubes. Serial dilutions of tissue samples were plated on brain heart infusion agar plates for 24 hours at 37 C’. Vaccinia viral titer and Listeria burden in tissues were normalized by the weight of the tissues.

### *In vivo* KLF2-GFP detection and DC-T cell priming

To assess *in vivo* KLF2-GFP expression, cDCs were analyzed in draining lymph nodes 24 hours after footpad injection with 25ul of the vaccine (containing 10ug chicken OVA with or without 10% ADJ or 10ug LPS). For *in vivo* DC-T cell priming, BMDCs were loaded with 1mg of OVA with or without 1% ADJ for 6 hours, washed twice with PBS, and injected intravenously into mice. Spleens were collected on day 7 and the percentages of CD44^HI^ H-2K^b^/SIINFEKL tetramer-binding CD8 T cells were quantified by flow cytometry.

### B3Z activation assay for *in vitro* cross presentation

The cross-presentation capacity of murine BMDCs was measured using B3Z hybridoma cells, as previously described (Ghosh and Shapiro, 2012, Karttunen et al., 1992). Briefly, DCs were plated at 1 x 10^5^ cells/well in 96-well round bottom culture-treated plate (Corning). In some experiments, BMDCs were pre-treated for 1 hour with aponocyin (300µM) leupeptin (50μM), α-tocopherol (100μM), epoxomicin (10μM) or xanthohumol (30μM). Subsequently, BMDCs were cultured with OVA (1mg/ml) with or without appropriate chemicals for 5 hours. Next, BMDCs were fixed with 0.025% glutaraldehyde for 2 minutes at room temperature. washed with PBS and cultured with B3Z cell (1 x 10^5^ cells/well) for 18 hours. After 18 hours, B3Z cells were washed and incubated with CPRG substrate (0.15mM) in 200ul of lysis buffer (0.1% NP 40+ PBS) for 18 hours at room temperature. The absorbance at 590nm was measured using a plate reader. Wells containing B3Z cells + BMDCs without OVA served as background control.

### Cytokine detection from cell culture supernatants

Supernatants from cultures of BMDCs treated with or without ADJ (1%) were collected at 24 hours. Cytokines from cell culture supernatants were quantified using Bio-Plex Pro Mouse Cytokine 23-plex Assay (Bio-rad), IFN-β ELISA kit (R&D Systems, DY8234-05), and IL-18 ELISA (Thermo Fisher Scientific, BMS618-3) according to manufacturer’s protocol.

### Antigen capture and processing assay

BMDCs were seeded at 4-5 x 10^5^ cells per well in 96-well culture flat-bottom plate (Corning) for the following assays. For antigen uptake assay, 20ug Alexa Fluor 647-conjugated OVA was mixed with or without ADJ (1%) in 200ul of complete RPMI media. BMDCs were incubated with pre-warmed AF-647 OVA with or without 1% ADJ for 10 or 30 minutes. The pH of OVA-containing endosomal compartment was determined using ovalbumin-conjugated with pHAB Amine Reactive dye as previously described with some modifications (Kar et al., 2016, Nath et al., 2016). Conjugation of ovalbumin with pHAB-amine reactive dye was performed according to manufacturer’s instructions. Briefly, OVA was labeled with amine-reactive dye at 1:10 excess molar ratio (ovalbumin:pHAB amine reactive dye). Free-unlabeled dye was removed by PD-10 columns (GE healthcare) and concentrated using Amicon® Ultra Centrifugal Filters (Millipore). BMDCs were incubated with pHAB-conjugated OVA with or without 1% ADJ for 30 minutes, and chased for 30 minutes. Antigen degradation studies were performed using DQ-OVA; DCs were incubated with DQ-OVA with or without 1% ADJ for 30 minutes, and chased for 6 hours.

### Detection of intracellular ROS, H_2_O_2,_ lipid peroxidation, and intracellular acidity

ROS and H_2_O_2_ production were measured using CellROX Deep Red reagent and ROS-Glo H_2_O_2_ assay, respectively. For intracellular H_2_O_2_ detection, ROS-Glo H_2_O_2_ assay was performed according to manufacturer’s instructions; luciferin activity from added H_2_O_2_ substrate was measured by SpectraMax i3x (Molecular Probes). For ROS production, samples were washed with PBS and incubated with 50nM CellROX Deep Red in PBS for 15 minutes at 37C. Lipid peroxidation was measured using Image-iT lipid peroxidation kit according to manufacturer’s instructions. For tracking acidic organelles, samples were washed and incubated with 50nM Lysotracker Deep Red in PBS for 15 minutes at 37C.

### Endosomal antigen leakage assay

Cytochrome C release assay was performed as previously described with brief modifications (Lin et al., 2008). Briefly, 5mg/ml of equine cytochrome C (Sigma Aldrich) was incubated with or without 1% ADJ in DCs. After 24 hours, cells were stained with Annexin-V and analyzed by flow cytometry.

### 20S proteasome activity assays

The activities of proteasomes containing luminogenic substrates (the Suc-LLVY, Z-LRR and Z-nLPnLD sequence recognized by the 20S proteasome) were measured by Proteasome-Glo™ Chymotrypsin-Like, Trypsin-Like and Caspase-Like Cell-Based Assays (Promega), according to manufacturer’s instructions (Moravec et al., 2009).

### Immunofluorescence, confocal laser scanning microscopy for co-localization

For co-localization of antigen-containing compartment, DCs were seeded in a 24-well plate at a density of 1× 10^6^ cells/well on 1.5mm coverslips (Warner Instrument) and cultured in complete phenol-red free RPMI 1640 media (Lonza) for 30 minutes at room temperature. DCs were pulsed with Alexa Fluor 647-OVA (40 ug/ml) +/-ADJ (1%) for 15 minutes at 37 C, and chased in complete phenol-red free RPMI media for 20 or 60 minutes at 37 C. Cells were washed with PBS, fixed with 4% paraformaldehyde (Electron Microscopy Services) at room temperature for 20 minutes, permeabilized using 0.2% Triton X-100 and blocked in blocking buffer (10% goat serum, 0.1% cold-fish gelatin, 0.1% tween-20) for 1 hour. Following antibodies that were diluted in blocking buffer were used to detect early endosomes and lysosomes at 4C’ overnight: rabbit anti-EEA1 (Clone: C45B10, 3288S, 1:50, Cell Signaling technology), rat anti-LAMP1 (Clone: 1D4B, sc-19992, 1:50, Santa Cruz Biotechnology), Coverslips were extensively washed with PBS + 0.1% Tween-20 and incubated with Alexa 564 conjugated rat IgG antibody and Alexa 488 conjugated rabbit IgG antibody (Invitrogen) in blocking buffer at room temperature for 1 hour. Samples were counter-stained with DAPI (Invitrogen), washed, mounted with Prolong Gold Anti-fade mountant (Invitrogen) and imaged on a Leica SP8 confocal laser-scanning microscope at 63X objective lens. The degree of co-localization was calculated using LAX-S Software (Leica).

### Metabolism assays

For real-time analysis of ECAR and OCR, Glycolysis Stress and Mito Stress tests were performed according to manufacturer’s instructions (Pelgrom et al., 2016). BMDCs (80,000 cells/well) were analyzed with an XF-96 Extracellular Flux Analyzer (Seahorse Bioscience). 1-2 x 10^5^ cells/well were used in 96-well tissue-culture treated flat bottom plates for the metabolic measurements; ATP concentrations, glucose, lactate, NAD+/NADH, and NADP+/NADPH were quantified using CellTiter-Glo®, Glucose-Glo, Lactate-Glo, NAD+/NADH-Glo, and NADP+/NADPH-Glo kits (Promega), respectively, according to manufacturer’s instructions. For accurate quantification of metabolites from the cell culture supernatant, dialyzed FBS (Gibco) was used in complete RPMI media. Nitrite levels were quantified using Griess’ reagent (Sigma Aldrich). 3-4 x 10^5^ cells/well were used in 96-well non-treated flat-bottom plates for the following metabolism assays and transferred to 96-well round-bottom plates for staining; mitochondrial contents, membrane potentials, and mitochondrial ROS were measured by treating cells with Mitotracker Deep Red, DiOC_6_, and MitoSOX, respectively in PBS for 30 minutes according to manufacturer’s protocol. To visualize neutral lipids, cells were stained with 500 ng/ml BODIPY 493/503 in PBS for 15 minutes; stained cells were acquired immediately on a flow cytometer without fixation.

### KLF2-GFP and ODD-luc reporter assay

BMDCs derived from ODD-Luc or KLF2-GFP mice were treated with ADJ (1%) or LPS (100ng/ml) For KLF2-GFP reporter experiments, samples were washed and immediately acquired on flow cytometer. For ODD-luc DCs, cells were lysed using passive lysis buffer and luciferase activities were measured using Luciferase® reporter assay system (Promega) according to manufacturer’s instructions.

### Quantification of the optical redox ratio by live cell microscopy

Optical imaging of BMDCs was performed as described previously (Walsh et al., 2013). BMDCs were seeded at a density of 1 x 10^6^/ ml in glass bottom dishes 2 hours before imaging. For the time course experiments, 1% ADJ was added immediately before imaging. Cells were imaged in a stage-top incubator maintained at 37 °C with CO_2_ supplementation. Four locations within each dish were imaged for control and ADJ-treated groups. The optical redox ratio was calculated from intensity images, which were acquired for 60 seconds as previously described (Walsh et al., 2013). The total number of NADH photons was divided by the sum of the total number of NADH photons and the total number of FAD photons on a single pixel basis. A custom Cell Profiler pipeline was used to threshold out nuclear signal and background. The average redox ratio value per cytoplasm was then computed.

### Immunoblot

Unstimulated or BMDCs stimulated with ADJ (1%) or LPS (100ng/ml) were washed with cold PBS and lysed in RIPA buffer (CST) containing protease and phosphatase inhibitor cocktail. Total protein levels in each lysate were estimated using Pierce BCA protein assay kit (Thermo Scientific). 30-40ug samples were loaded and resolved on a 12% SDS-PAGE gel (Genscript), transferred to PVDF membrane using iBlot 2 Gel Transfer (Thermo Fisher Scientific), blocked with 5% BSA in TBST for phospho-proteins and 5% milk for total proteins in TBST for 1 hour and probed with primary antibodies (at 4C overnight). Primary rabbit antibodies were used as follows at 1:1000 dilution in 5% BSA in TBST: Rabbit monoclonal anti-phospho-Akt (Ser473; Clone: D9E), Rabbit polyclonal anti-total Akt, Rabbit monoclonal anti-phosphor-p70 S6 kinase (Thr389, Clone: 108D2), and Rabbit polyclonal anti-p70 S6 kinase antibodies. Blots were extensively washed with TBST and incubated with goat anti-rabbit IgG (H+L)-HRP antibodies (Thermofisher) diluted in 5% non-fat milk in TBST for 1 hour at room temperature. Protein bands were visualized by ECL prime western blotting detection reagent (GE Healthcare.) The membranes were stripped with mild stripping buffer (0.1% SDS, 0.1% Tween 20, 1.5% Glycine), when necessary.

### Discovery lipidomics by LC-MS

Samples were spun down and snap frozen in liquid N_2_ and stored at −80C until extraction. Media samples were thawed on ice, and 50ul of media was transferred to microcentrifuge tube; cell pellets were directly extracted in tubes. To perform extraction, 187.5μL of chilled methanol and 750μL of chilled methyl tert-butyl ether (MTBE) was added to each tube, followed by the addition of a 5mm stainless steel bead. All samples were then bead-beaten in a cold room on a Retsch MM400 mixer mill at a frequency of 25 cycles/sec for 5 minutes to complete cell lysis. This mixture was then vortexed for 1-2 s and quickly centrifuged to remove any stray droplets from the tube openings. Then 187.5μL of chilled water was added to each tube to separate the hydrophobic and hydrophilic compounds into separate phases. All samples were then vortexed for 30 seconds and then centrifuged at 4C for 2 minutes at a speed of 14000 x g to re-pellet any cell debris. A total of 300μL of the top-layer of the biphasic extraction was removed from the tube and collected into a low volume borosilicate glass autosampler vial with tapered insert and dried by vacuum concentrator. All samples were reconstituted with 50μL of a 9:1 MeOH:Toluene solution for injection.

LC-MS analysis was performed on an Acquity CSH C18 column held at 50 °C (100 mm x 2.1 mm x 1.7 μm particle size; Waters) using a Vanquish Binary Pump (400μL/min flow rate; Thermo Scientific). Mobile phase A consisted of 10mM ammonium acetate and 250μL/L acetic acid in ACN:H2O (70:30, v/v). Mobile phase B consisted of IPA:ACN (90:10, v/v) with the same additives. Initially, mobile phase B was held at 2% for 2 min and then the following gradient was employed: increase to 30% over 3 min, then to 50% over 1 min, then to 85% over 14 min, and finally to 99% over 1 min where %B was held at 99% for 7 min. The column was then re-equilibrated with mobile phase B at 2% for 1.75 min before the next injection. 10μL of each extract was injected by a Vanquish Split Sampler HT autosampler (Thermo Scientific) in a randomized order. The LC system was coupled to a Q Exactive HF Orbitrap mass spectrometer (MS) through a heated electrospray ionization (HESI II) source (Thermo Scientific). Source conditions were as follows: HESI II and capillary temperature at 350C, sheath gas flow rate at 25 units, aux gas flow rate at 15 units, sweep gas flow rate at 5 units, spray voltage at |3.5 kV|, and S-lens RF at 90.0 units. The MS was operated in a polarity switching mode acquiring positive and negative full MS and MS2 spectra (Top2) within the same injection. Acquisition parameters for full MS scans in both modes were: 30,000 resolution, 1 × 106 automatic gain control (AGC) target, 100 ms ion accumulation time (max IT), and 200 to 2000 m/z scan range. Data dependent (dd-MS2) scans in both modes were then collected at 30,000 resolution, 1 × 105 AGC target, 50 ms max IT, 1.0 m/z isolation window, stepped normalized collision energy (NCE) at 20, 30, 40, with a 10.0 s dynamic exclusion. The resulting LC–MS/MS data were processed using Compound Discoverer 2.1 (Thermo Scientific) and LipiDex (PMID: 29705063), an in-house-developed software suite. All peaks with a 0.2 min to 23 min retention time and 100 Da to 5000 Da MS1 precursor mass were aggregated into distinct chromatographic profiles (i.e., compound groups) using a 10-ppm mass and 0.5 min retention time tolerance. Profiles not reaching a minimum peak intensity of 1×106, a maximum peak-width of 0.35, a signal-to-noise (S/N) ratio of 3, and a 3-fold intensity increase over blanks were excluded from further processing. MS/MS spectra were searched against an in-silico generated lipid spectral library containing 35,000 unique molecular compositions representing 48 distinct lipid classes. Spectral matches with a dot product score greater than 500 and a reverse dot product score greater than 700 were retained for further analysis. Lipid MS/MS spectra which contained no significant interference (<75%) from co-eluting isobaric lipids, eluted within a 3.5median absolute retention time deviation (M.A.D. RT) of each other, and found within at least 2 processed files were then identified at the individual fatty acid substituent level of structural resolution. If individual fatty acid substituents were unresolved, then identifications were made with the sum of the fatty acid substituents. Lipid quantitation was normalized to cell numbers.

### Data availability

Discovery lipidomics raw files and results tables can be accessed through MassIVE [doi:10.25345/C52X26], repository identifier MSV000085222. For reviewers, these data can be accessed with username: MSV000085222_reviewer, password: view_data.

## Quantification and Statistics

All experiments are performed and repeated 2-5 times; data are representative of 2-5 independent experiments. Data are presented as the mean ± SEM. Student’s two-tailed t-test, and one-way ANOVA analyses were used to calculate the statistical significance of differences between groups, and significance was defined at p < 0.05. Statistical Differences in measured variables between the experimental and control groups were assessed using Student’s t test and p < 0.05 was considered as statistically significant. Stars are p values in the following ranges: 0 - 0.001 = ‘***’, 0.01 - 0.05 = ‘*’, 0.05 - 0.1 = ‘.’.

## Acknowledgements

We would like to thank all the members of Suresh Laboratory for constructive feedbacks and technical assistance. Special thanks to Drs. Natalie Niemi and David Pagliarini for technical assistance for the use of Seahorse Bioanalyzer. Thanks to Dr. Sathish Kumar for the use of SpectraMax i3x plate reader. We appreciate help by Drs. Gregory Wipez and Gopal Iyer for Bioplex assay and immunofluorescence; Thanks to Zachary Morrow for Listeria challenge experiment. We are thankful to the Emory NIH Tetramer Core Facility for providing MHC-I tetramers.

## Author contributions

W.L, B.B, B.P, A. L, K. D, C.M, R.T, G.C, K. M, M.S. designed, performed, analyzed experiments. K.T, K.O, A.W, J. R provided technical analysis and critical expertise. W.L., K.S, L.R, M.S, J.C, K.R, M.S. provided conceptual input for the manuscript. J.S, D.S provided reagents critical for this manuscript. W.L and M.S wrote the manuscript, which was proofread by all authors.

---

**Supplementary Figure 1:**
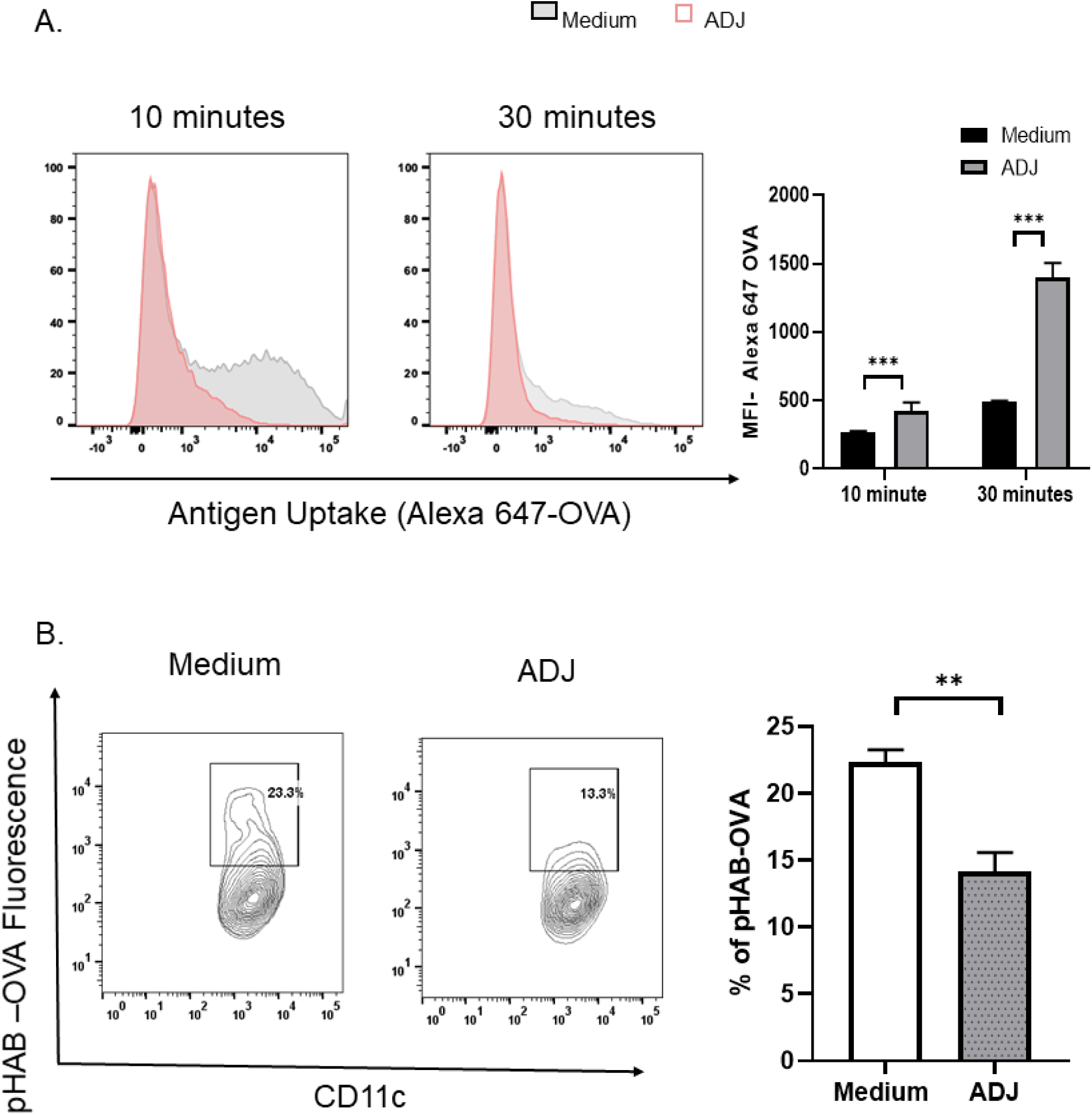
Carbomer-based adjuvant alters antigen uptake and processing in DCs. (A) Kinetics of antigen uptake by BMDCs treated with ADJ. Cells were cultured with 20ug/ml OVA-Alexa Fluor 647 (pH insensitive dye) with or without 1% ADJ for 10 and 30 minutes. (B) Effects of ADJ on intracellular routing of antigens. BMDCs were cultured with 20ug/ml OVA labeled with the pH sensitive dye (pHAB), with or without 1% ADJ for 30 minutes. Data are representative of ≥2 independent experiments Error bars are SEM; \**P*<0.01; \*\**P*<0.001; \*\*\**P*<0.0001 (Student’s t-test and one-way ANOVA).

**Supplementary Figure 2.**
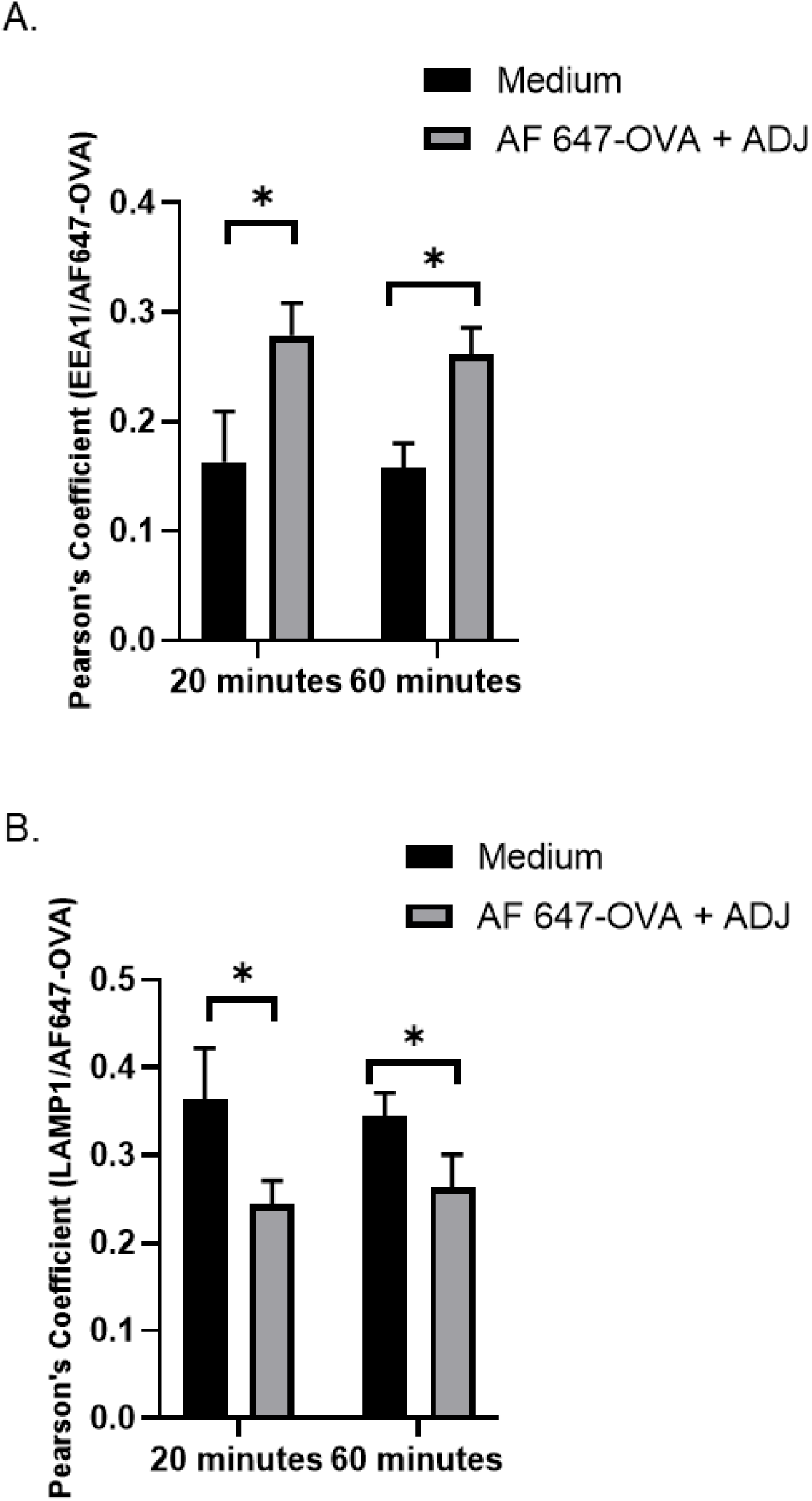
Carbomer-based adjuvant promotes intracellular routing of OVA to early endosomes. BMDCs were pulsed with Alexa Fluor 647-OVA (60 µg/ml) and chased at the indicated time-points to assess EEA1 (A) or LAMP1 (B) co-localization. Pearson’s coefficient was calculated from 10 cells/treatment. Data are representative of ≥2 independent experiments Error bars are SEM; \**P*<0.01; \*\**P*<0.001; \*\*\**P*<0.0001 (Student’s t-test and one-way ANOVA).

**Supplementary Figure 3.**
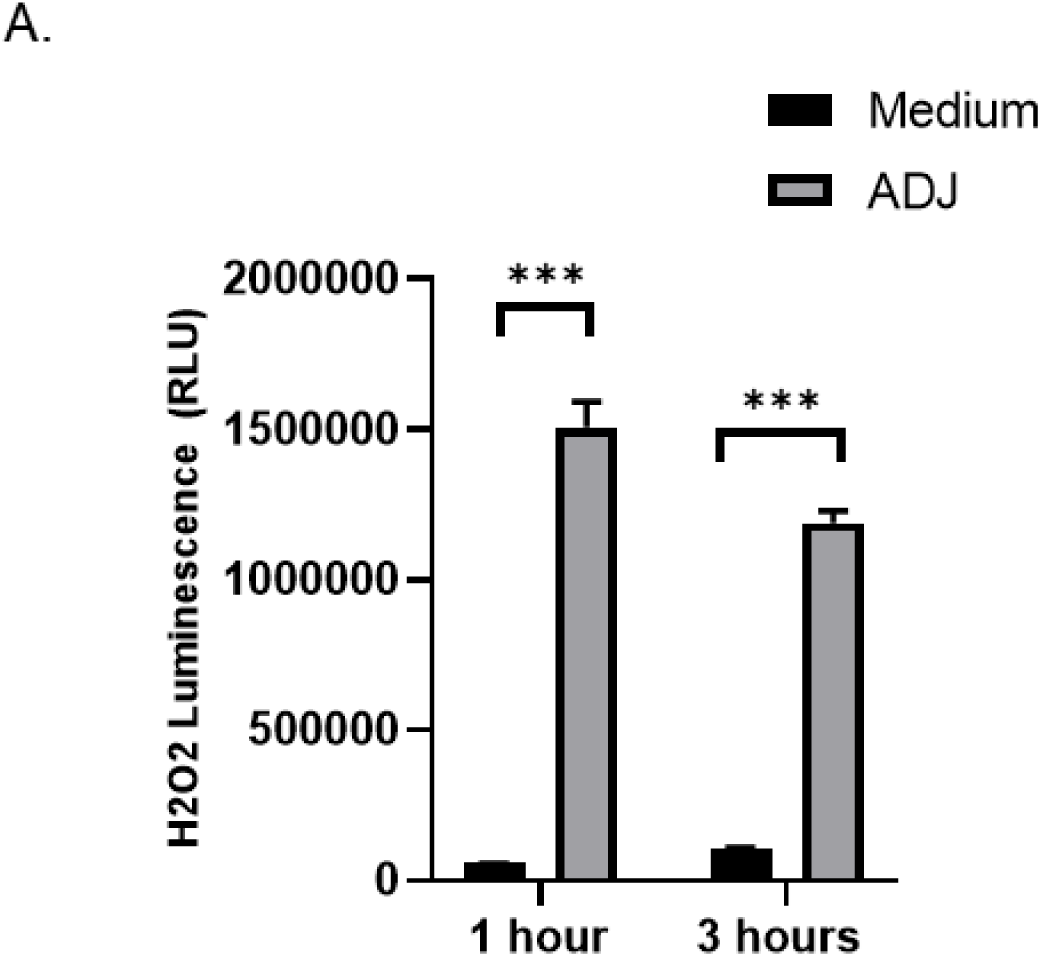
Carbomer-based adjuvant induces intracellular H_2_O_2_ production in DCs. BMDCs were treated with 1% ADJ for 1 and 3 h with luminogenic substrate. H_2_O_2_ levels were detected by ROS-Glo™ detection solution. Data are representative of ≥2 independent experiments Error bars are SEM; \**P*<0.01; \*\**P*<0.001; \*\*\**P*<0.0001 (Student’s t-test and one-way ANOVA).

**Supplementary Figure 4.**
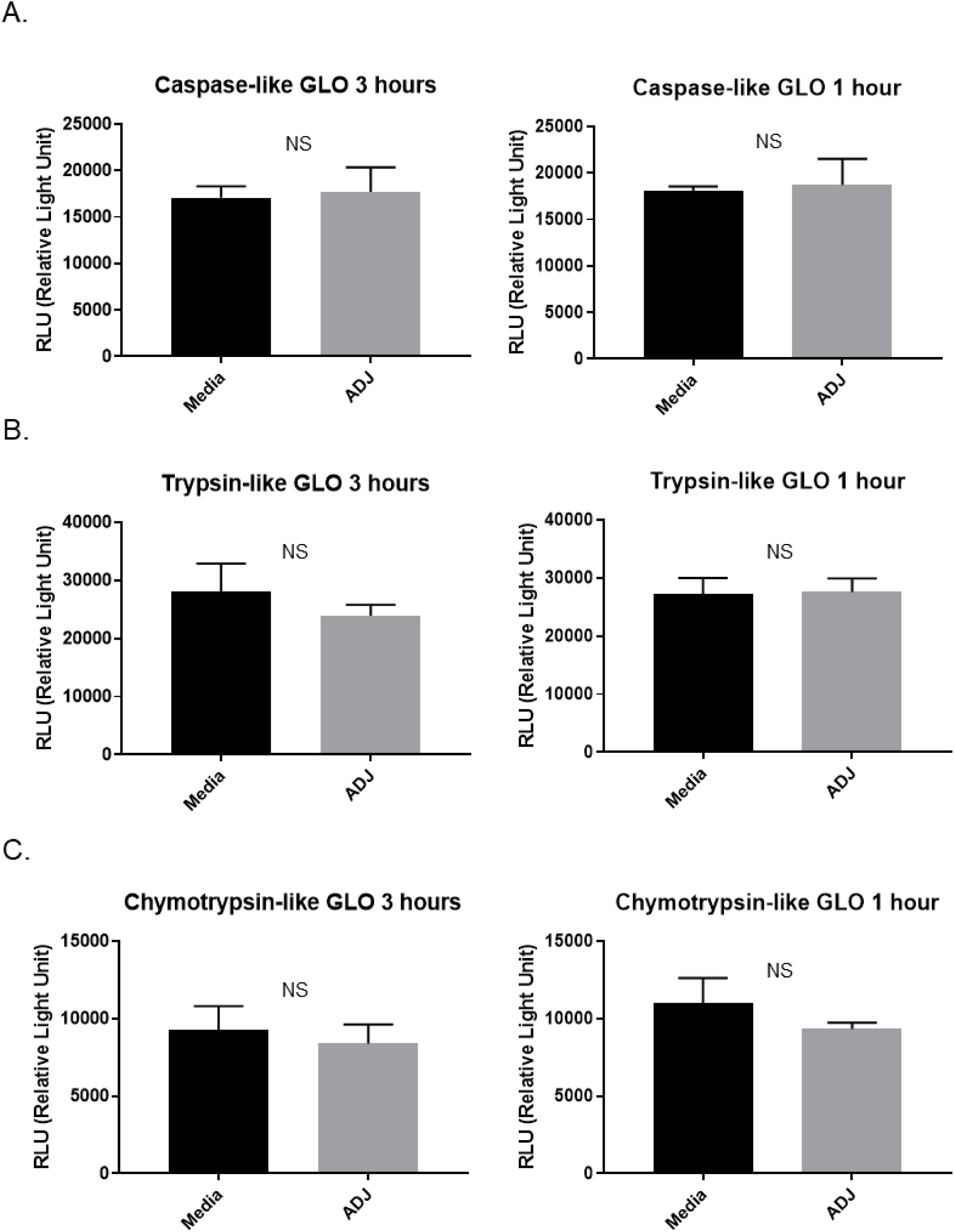
Carbomer-based adjuvants do not affect proteasome activities in DCs. BMDCs were stimulated with 1% ADJ for 1 or 3 h and incubated with specific luminogenic proteasome substrates Suc-LLVY (A), Z-LRR (B), and Z-nLPnLD (C) for the chymotrypsin-like, trypsin-like and caspase-like activities, respectively. Following cleavage by the proteasome, the substrate for luciferase is released and the luminescence was detected using plate reader. Data are representative of ≥2 independent experiments Error bars are SEM; \**P*<0.01; \*\**P*<0.001; \*\*\**P*<0.0001 (Student’s t-test and one-way ANOVA).

**Supplementary Figure 5.**
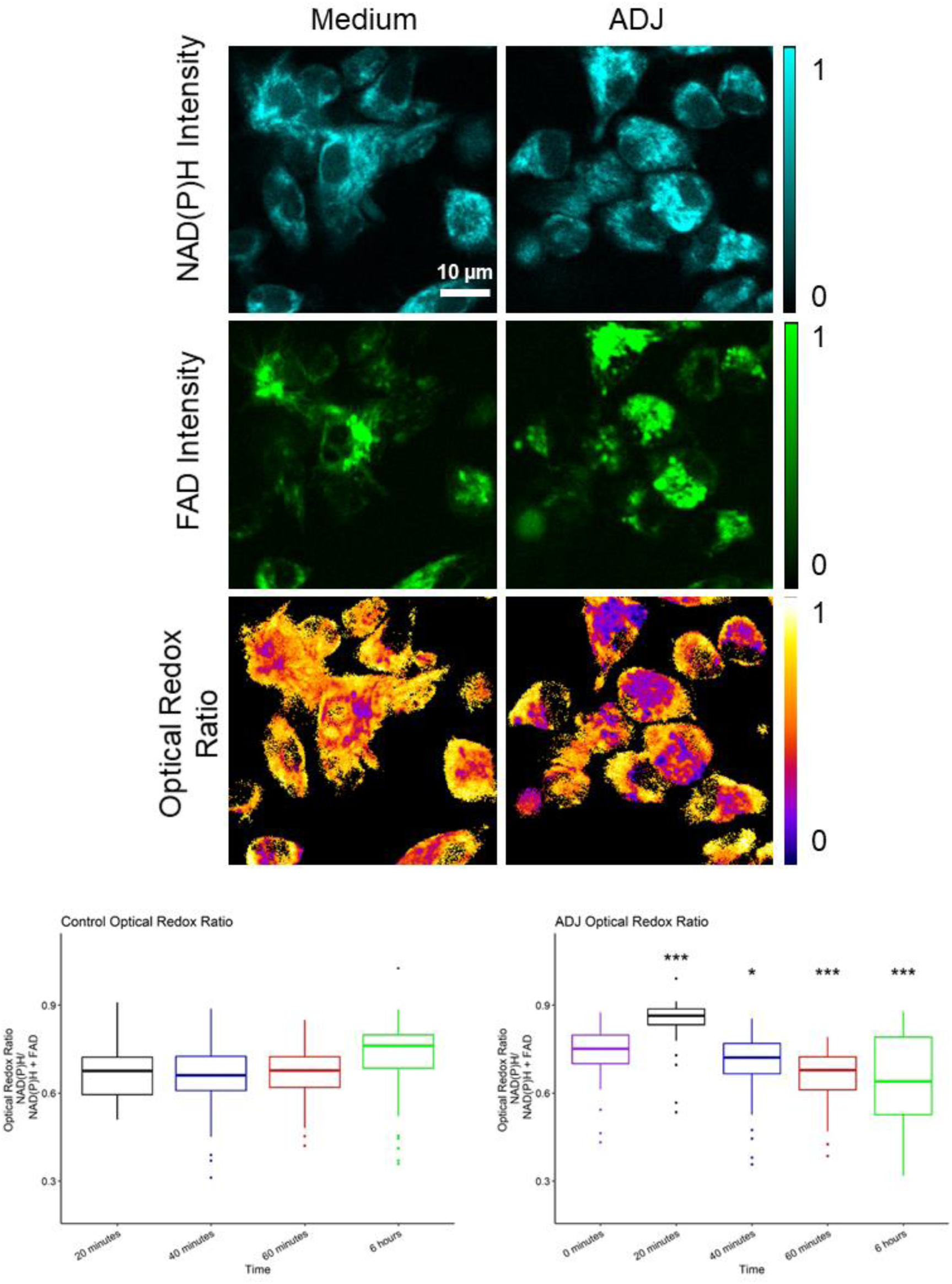
Carbomer-based adjuvants reduce optical redox ratio in DCs. Optical redox ratio of unstimulated and ADJ treated dendritic cells was calculated at indicated time. Representative NAD(P)H intensity (first row), FAD intensity (second row), and optical redox ratio (NAD(P)H/(NAD(P)H+FAD); third row) images of unstimulated and ADJ-treated dendritic cells. Scale bar is 10 µm. Box plots show median (central line), first and third quartiles (lower and upper hinges), the farthest data points that are no further than 1.5* the interquartile range (whiskers), and data points beyond 1.5* the interquartile range from the hinge (dots). Stars compare respective boxes to the first time point of each group (n=22-75 cells/time point). Data are representative of ≥2 independent experiments with ≥3 mice or triplicates per group. Error bars are SEM; \**P*<0.01; \*\**P*<0.001; \*\*\**P*<0.0001 (Student’s t-test and one-way ANOVA).

